# Analyzing the role of a dual-function diguanylate cyclase/phosphodiesterase for attachment and motility in *Agrobacterium tumefaciens*

**DOI:** 10.1101/2025.08.14.670421

**Authors:** Priya Aryal, Matthew Anderson, Hannah Turrisi, Aathmaja Anandhi Rangarajan, Mackenzie Brooks, Gianna Rodriguez, Delayna Warrell, Anthony Pangalinan, Ganishah Charles-Estain, Ravinder Abraham, Hope Zaborowski, Dalal Seyhan, Christopher M. Waters, Jason E. Heindl

## Abstract

Cyclic diguanylate monophosphate (c-di-GMP) is a second messenger that controls the signaling pathway for the motile to sessile transition in many bacteria. Diguanylate cyclases (DGCs), characterized by a GGDEF domain, are responsible for the synthesis of c-di-GMP. c-di-GMP is degraded by EAL and HD-GYP domains that exhibit phosphodiesterase activity (PDE). In *Agrobacterium tumefaciens*, out of 31 predicted proteins regulating the c-di-GMP levels, this work focuses on the predicted dual-function DGC/PDE, DcpB. We show that DcpB is a cycle-dependent PDE under our experimental conditions, resulting in cell cycle-dependent control of motility and biofilm formation in *A. tumefaciens*. DcpB also exhibits polar and mid-cell localization. Finally, we identify genetic interactions between *dcpB* and a LacI-family transcriptional regulators, *thuR*, suggesting additional regulatory inputs for DcpB-dependent phenotypes.

**Importance:** c-di-GMP is a universal secondary messenger known to control various biological processes in bacteria including attachment, motility, virulence, and cell cycle progression. In *A. tumefaciens*, there are 31 predicted c-di-GMP-metabolizing proteins predicted to affect the c-di-GMP pool, raising the question of why so many proteins are involved and how the activities of these proteins are coordinated. This work describes a cell cycle-dependent PDE activity that helps coordinate cell cycle progression and developmental phenotypes including biofilm formation and motility.

## Introduction

Cyclic diguanylate monophosphate (c-di-GMP) is an intracellular second messenger that controls the signaling pathway for the motile to sessile transition in many bacteria (1, 2). In many studied bacteria, c-di-GMP has a reciprocal effect on attachment and motility. Increased levels of c-di-GMP positively regulate attachment resulting in biofilm formation and negatively regulate motility (1, 2). c-di-GMP is metabolized by the opposing activities of diguanylate cyclase enzymes (DGC) and cyclic-di-GMP-specific phosphodiesterase enzymes (PDE). DGCs are characterized by a c-di-GMP-synthesizing domain, termed DGC or GGDEF domain, which contains a canonical GGDEF amino acid motif in the active site. PDEs are characterized by one of two recognized c-di-GMP-degradative domains, generically termed PDE domain and more specifically annotated as either an EAL or HD-GYP domain, with reference to the conserved EAL or HD-GYP amino acid residues, respectively. It is not unusual for a particular bacterial species to have multiple DGCs, PDEs, or proteins containing both domains, so-called dual-domain DGC/PDE proteins.

In addition to regulation by c-di-GMP, differentiation between motile, planktonic cells and non-motile, attached cells may be an intrinsic property of the bacterial cell cycle. For many bacteria, cells transition between two distinct cell types following daughter cell separation. This biphasic cell cycle is well-documented among members of the alphaproteobacterial class, notably within the orders *Caulobacterales* and *Hyphomicrobiales*. The *C. crescentus* cell cycle is regulated by the PdhS-DivK-CtrA pathway. In this pathway, a set of sensor histidine kinases, PleC and DivJ, founding members of the PdhS family of sensor histidine kinases, controls the phosphorylation status of the receiver domains of the DivK and PleD response regulators. DivK and PleD, in turn, ultimately control the activity of the global transcriptional regulator, CtrA. CtrA,, directly and indirectly controls transcription of hundreds of target genes. CtrA activity is itself regulated by receiver domain phosphorylation, controlled by the bifunctional hybrid histidine kinase, CckA. CckA is biased towards phosphatase activity when DivK is phosphorylated. Additionally, c-di-GMP allosterically biases CckA towards phosphatase activity (3).

In *C. crescentus*, cell cycle regulation and the motile-to-sessile switch intersect due to a combination of c-di-GMP-specific effectors and alteration of levels of c-di-GMP (4). The response regulator, PleD, contains a DGC domain which is activated upon phosphorylation of its receiver domain. Active, phosphorylated PleD localizes to the stalked cell pole of the bacterium resulting in increased local concentrations of c-di-GMP at this location (5). These increased levels of c-di-GMP at the stalked cell pole contribute to decreased CtrA activity by interacting with CckA as described above, and by facilitating ClpXP-mediated regulated proteolysis via the PopA adapter protein (3, 6, 7) . PopA itself is an enzymatically inactive diguanylate cyclase which instead binds c-di-GMP, functioning as a c-di-GMP-dependent effector protein. A second diguanylate cyclase, DgcB, is a primary contributor to high levels of c-di-GMP in the stalked cell compartment of *C. crescentus*, while in the swarmer cell compartment c-di-GMP levels are depleted by the activity of the dual-domain diguanylate cyclase/phosphodiesterase, PdeA (8).

*Agrobacterium tumefaciens* is a plant pathogen from class *Alphaproteobacteria* known to cause crown gall disease by transforming host plant cells via type IV secretion system-mediated transfer and subsequent integration of a segment of its T-DNA, into the host cell (9, 10). *A. tumefaciens* is known to form a biofilm on both abiotic and biotic surfaces and exhibits flagellum-dependent swimming motility (11, 12) One of the factors affecting attachment and motility in *A. tumefaciens* is c-di-GMP. In *A. tumefaciens* there are 31 predicted proteins regulating the intracellular level of c-di-GMP: 16 GGDEF-containing DGCs, 1 EAL-only PDE, 1 HD-GYP-only PDE, and 13 GGDEF-EAL dual-domain DGC/PDEs (1, 13). Of these, several have been experimentally determined to contribute to the c-di-GMP pools regulating attachment and motility phenotypes. These include the *A. tumefaciens* homologue of PleD (Atu1279) and a protein with similar architecture, Atu1060; three DGC enzymes identified as a part of the VisR regulon, DgcA (Atu1257), DgcB (Atu1691), and DgcC (Atu2179); and a dual-domain DGC/PDE, DcpA (Atu3495) whose activity is controlled by a recently described novel tetrahydromonapterin-binding protein, PruR (Atu3496) (12, 14, 15). The remaining predicted c-di-GMP-metabolizing proteins have not been explored well in *A. tumefaciens* and understanding their effects on c-di-GMP-dependent phenotypes is an important area of current study. Cell cycle regulation in *A. tumefaciens* uses the conserved PdhS-DivK-CtrA regulatory pathway (16–18). Overall PdhS-DivK-CtrA pathway architecture is conserved between *C. crescentus* and *A. tumefaciens*, with primary input being a suite of sensor histidine kinase homologues related to PleC and DivJ: PleC (Atu0982), DivJ (Atu0921), PdhS1 (Atu0614), and PdhS2 (Atu1888), and primary outputs being transcriptional regulation by CtrA (Atu2434) and c-di-GMP production by PleD (18). How c-di-GMP levels are cleared in motile daughter cells following the asymmetric cell division of *A. tumefaciens* remains unclear.

Here, we identify DcpB (Atu3207) as a cell cycle-dependent PDE responsible for eliminating high levels of c-di-GMP in *A. tumefaciens*. DcpB is a predicted dual-function DGC/PDE containing six transmembrane helices and is associated with intact, and predicted enzymatically active, GGDEF and EAL domains. We show that DcpB primarily acts as a PDE, negatively regulating attachment and having a reciprocal effect on motility, and that these phenotypes depend on DcpB directly affecting intracellular levels of c-di-GMP. Moreover, our data support polar localization of DcpB and CtrA-dependent transcription of *dcpB*. Overall, our results suggest that DcpB affects c-di-GMP metabolism, resulting in cell cycle-dependent control of motility and biofilm formation in *A. tumefaciens*, most likely functioning as a cell cycle-dependent PDE under our experimental conditions.

## Results

### DcpB is a c-di-GMP phosphodiesterase

In the context of a course-based undergraduate research experience, we generated a transposon mutant library and screened for suppressors of the Δ*pdhS1* motility phenotype (data not shown) (18). Disruption of Atu3207 (designated as DcpB) was one of several transposon mutants isolated in this screen. Upstream of DcpB is another predicted DGC, Atu3204, and downstream of DcpB is one of two predicted FtsK paralogs, FtsK2 (Atu3210) (Fig. 1A). The predicted domain structure of DcpB consists of a conserved DGC domain with a GGDEF motif and an EAL domain with a PDE motif, along with six transmembrane helices (Fig. 1B).

**Fig 1.**
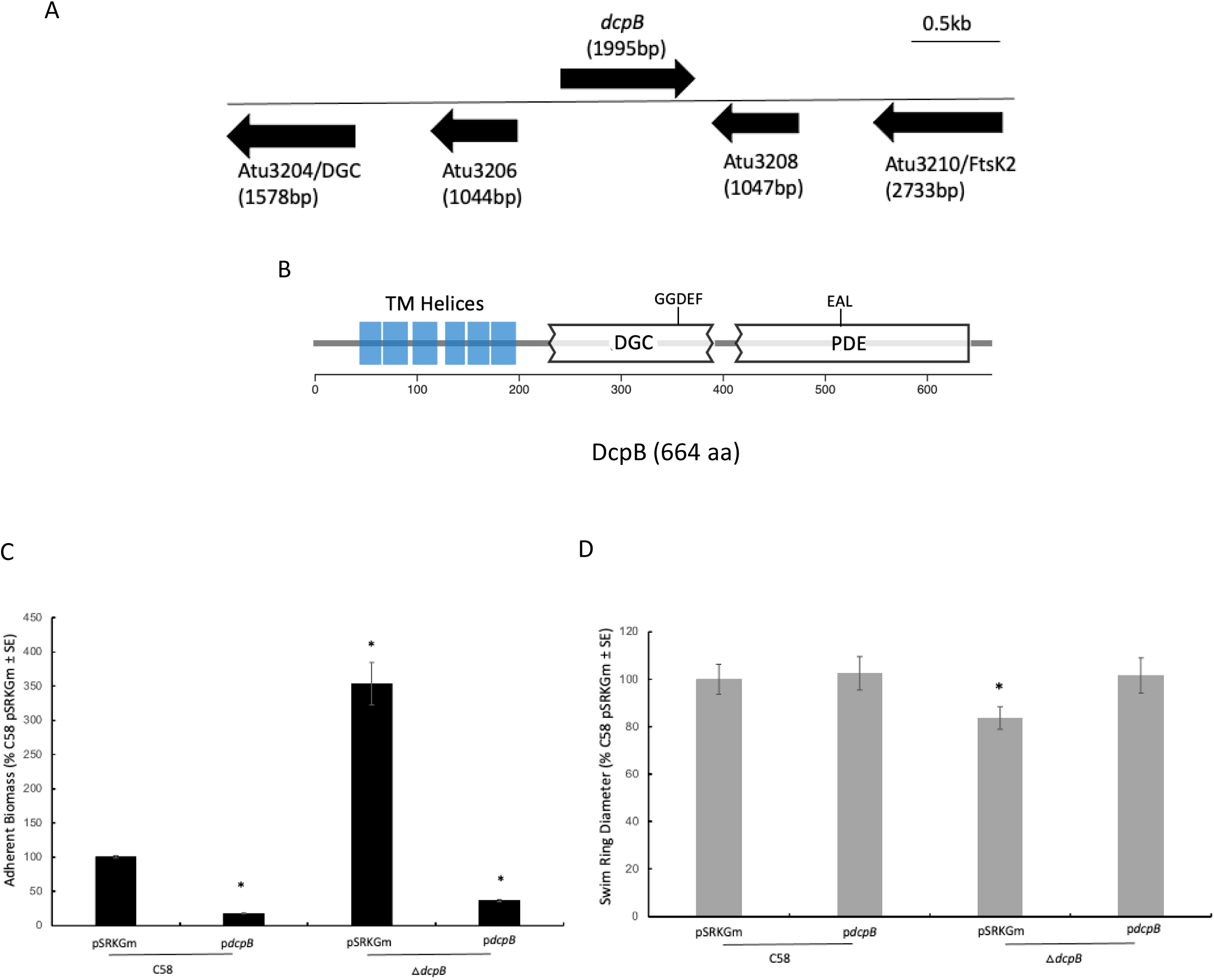
*dcpB* inversely regulates biofilm formation and swimming motility of *A. tumefaciens*. **A**. Chromosomal neighborhood of genetic locus *dcpB*. Upstream of *dcpB* is the predicted diguanylate cyclase Atu3204. Downstream genes include one of two predicted FtsK paralogs, Atu3210. **B.** Predicted domain structure of DcpB, a 664 amino acids-long protein with six transmembrane helices, one diguanylate cyclase domain with a conserved GGDEF motif, and one phosphodiesterase domain with a conserved EAL motif. TM, transmembrane; DGC, diguanylate cyclase; PDE, phosphodiesterase. Model generated using Microbial Signal Transduction Database 4.0 (https://mistdb.com/). **C.** 48 hours static biofilm assay was performed on wild-type and △*dcpB* strains carrying empty vector, pSRKGm, or plasmid borne *dcpB* (p*dcpB*). Adherent biomass was stained with crystal violet (CV). Optical density at 600 nm (OD_600_) measured culture growth, and absorbance at 600 nm measured solubilized CV. All strains were induced with 100 μM IPTG. Data were normalized to wild-type C58 carrying empty vector. ANOVA Single Factor with Tukey Kramer was used to determine statistical significance for n = 9. *, P < 0.05. **D.** A plate-based swim assay using low-density motility agar was performed on wild-type and △*dcpB* strains carrying empty vector, pSRKGm, or plasmid borne *dcpB* (p*dcpB*). Swim ring diameter was measured daily for seven days. Day seven data are shown. All strains were induced with 100 μM IPTG. Data were normalized to wild-type C58 carrying empty vector. Welch’s T-Test: Two-Tailed, Unequal Variance was used to determine statistical significance for n = 9 *, P < 0.05 compared to empty vector induced.

Ectopic expression of inducible, plasmid-borne DcpB (p*dcpB*) resulted in a significant reduction in biofilm formation relative to the wild-type C58 strain with empty vector (Fig.1C). Ectopic expression of DcpB and loss of *dcpB* expression showed reciprocal effects in these phenotypes with a robust increase in biofilm formation for the Δ*dcpB* strain (Fig. 1C) and a 20% reduction in swim ring diameter in this background relative to empty vector, suggesting low levels of global c-di-GMP in the wild-type C58 strain (Fig. 1D). Furthermore, p*dcpB* expressed in Δ*dcpB* strain (Δ*dcpB* p*dcpB*) was able to complement the phenotype for both biofilm formation and swimming motility (Fig. 1C, D). The reciprocal effect on biofilm and motility suggests regulation by c-di-GMP.

DcpB was originally identified during transposon mutagenesis in a Δ*pdhS1* background, where transposon insertion in *dcpB* suppressed the known reduced motility phenotype of the Δ*pdhS1* strain, increasing motility in this background (data not shown). To validate this, we tested the effect of DcpB in this background. Furthermore, to explore the effect of DcpB on motility in general, we chose to study the effect of increased DcpB activity in two additional backgrounds with known motility defects. The Δ*pdhS2* strain background, lacking another sensor kinase closely related to PdhS1, was selected due its integration with PdhS1 in the PdhS-DivK-CtrA pathway. The Δ*visR* strain background, a transcriptional regulator, was selected as a second, separate mutant background (12, 17). The Δ*pdhS1*, Δ*pdhS2,* and Δ*visR* strains carrying empty vector all maintain the biofilm formation and motility phenotypes as previously studied (Fig. S1 – 3). Expression of plasmid-borne DcpB in the Δ*pdhS1*, Δ*pdhS2,* and Δ*visR* strains reduced biofilm formation as was seen in the △*dcpB* mutant strain (Fig. S1A, S2A, S3A). In contrast, the motility phenotype was not affected when expressing p*dcpB* in any of these backgrounds (Fig. S1B, S2B, S3B). These results demonstrate that while the original transposon insertion in *dcpB* in the Δ*pdhS1* suggested DcpB acts as a c-di-GMP-specific cyclase, plasmid-borne wild-type DcpB is behaving as a c-di-GMP-specific phosphodiesterase.

To understand if DcpB is functioning as a c-di-GMP-specific cyclase or c-di-GMP-specific phosphodiesterase, or both, site-directed mutagenesis (SDM) was performed on each domain. Plasmid-borne ectopic expression constructs were generated where the GGEEF catalytic motif was mutated to GGAAF (pGGAAF), the EAL degradative motif was mutated to AAL (pAAL), or both motifs were mutated (pGGAAF-AAL). These mutations have been studied to inactivate the DGC and PDE domains (12, 15, 19–21). There was a significant reduction in adherent biomass when either pGGAAF or plasmid-borne p*dcpB* expression was induced with IPTG, compared to empty vector in both backgrounds (Fig. 2A). In contrast, no significant effect on adherent biomass was observed for either the pAAL or the dual domain mutation, pGGAAF-AAL, compared to the control (Fig. 2A). The modest increase in adherent biomass, relative to empty vector, when pAAL was expressed in the Δ*dcpB* mutant background suggests that there might be a functional GGEEF domain. We next tested swimming motility using low density (0.3%) swim agar plates. Significant increases in swim ring diameter were observed compared to the empty vector in the Δ*dcpB* mutant background when either plasmid-borne p*dcpB* or pGGAAF were induced (Fig. 2B). A similar phenotype as that of the empty vector control was observed for both pAAL and pGGAAF-AAL induction in the Δ*dcpB* mutant background. There was no significant difference in motility in the wild-type background (Fig. 2B).

**Fig 2.**
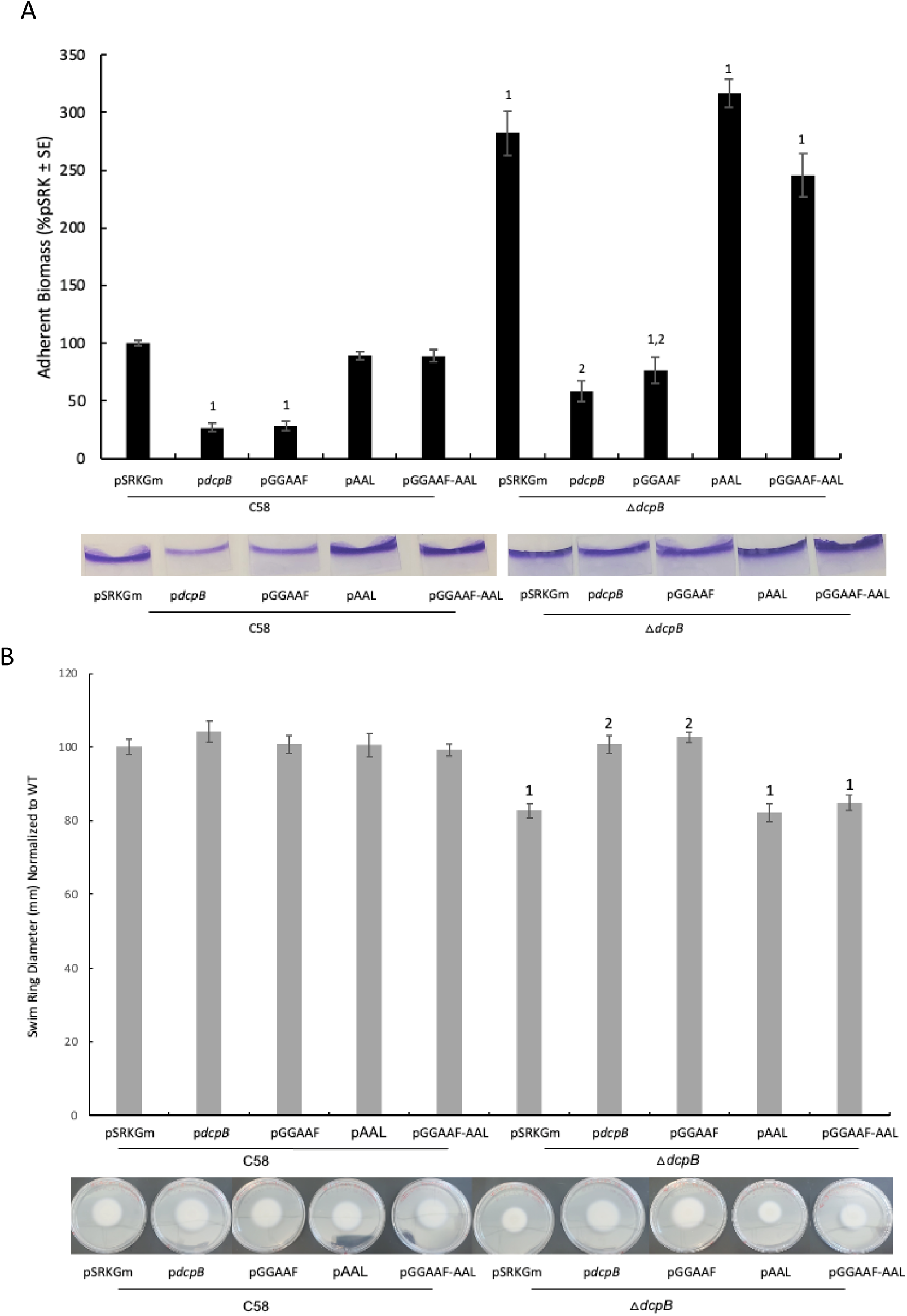
An intact EAL motif of *dcpB* is required for efficient complementation of observed △*dcpB* biofilm and motility phenotypes. **A**. 48 hours static biofilms were conducted as in Figure 1C on wild-type and △*dcpB* strains carrying the following plasmid-borne alleles of DcpB, as indicated: vector only (pSRKGm); wild-type *dcpB* (p*dcpB*); inactive GGDEF motif/intact EAL motif (pGGAAF); intact GGDEF/inactive EAL motif (pAAL); inactive GGDEF and AAL motifs (pGGAAF-AAL). Representative stained coverslips are included for each condition beneath the quantitative data. **B**. Plate-based swim ring assays were conducted as in Figure 1D on the same strains from Panel A. Day seven data are shown. Representative swim plates are included for each condition beneath the quantitative data. All strains were induced with 100 μM IPTG. Data were normalized to wild-type C58 carrying empty vector. Two-way ANOVA was used was used to determine statistical significance for n = 9. *, ^1^ P < 0.05 compared to wild-type C58 pSRKGm, ^2^ P < 0.05 compared to △*dcpB* pSRKGm.

*A. tumefaciens* can transfer its DNA (T-DNA) to the host leading to tumor formation by the host tissue. Tumor formation is another phenotype affected by c-di-GMP (22). A potato tumor assay was performed to understand the effect on virulence of DcpB. Compared to the C58 wild-type strain of *A. tumefaciens,* there was no significant change in tumor formation with the deletion of *dcpB* (Fig. S4). These data suggest that DcpB does not affect the tumor formation process. Overall, our biofilm and motility data suggest that the DcpB might be functioning as a c-di-GMP-specific phosphodiesterase.

### Loss of *dcpB* affects growth rate and cell size

Growth curves were conducted in ATGN media along with the complementation strain, Δ*dcpB* p*dcpB*. The Δ*dcpB* mutant strain’s growth rate was modestly reduced compared to C58 (Fig. 3A). However, the complementation strain induced with IPTG was able to restore the growth rate (Fig. 3A). We further studied the morphology of the Δ*dcpB* mutant strain. Although no obvious morphological difference was seen for the Δ*dcpB* mutant strain using phase contrast microscopy, we measured cell length using MicrobeJ, an ImageJ plugin. and found that d *dcpB* resulted in a smaller cell length compared to C58 wild-type (Fig. 3B).

**Fig 3.**
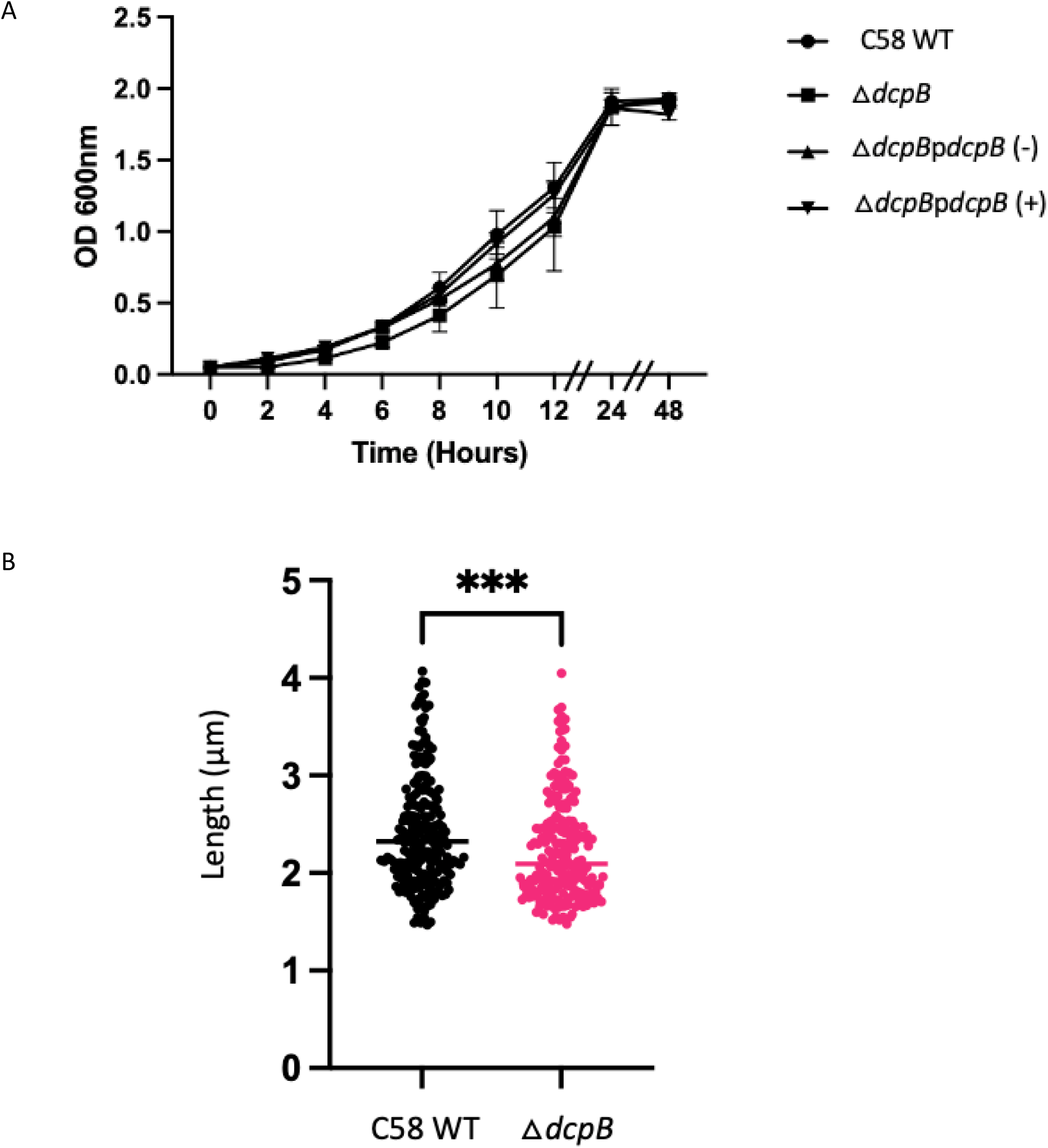
Absence of *dcpB* results in minimal affect on growth rate and resulted in smaller cell length by *A. tumefaciens*. **A.** Growth curve was constructed by measuring optical density at 600 nm (OD_600_) every two hours for the first 12 hours following sub-culture, and then at 24 hours and 48 hours. Overnight cultures grown in ATGN media were sub-cultured to OD_600_ = 0.05. Sub-cultures were, (+), or were not, (-), induced with 100 µM IPTG. No statistically significant differences are observed using a two-way ANOVA with Tukey-Kramer. P < 0.05. **B.** Phase contrast imaging of wild-type C58 and the △*dcpB* strain was collected using a Leica DMi8 inverted microscope. Data were analyzed using MicrobeJ. Sample size totaled 228 wild-type C58 cells and 217 △*dcpB* cells from two data sets with three fields of view. Statistical significance was determined using an Unpaired T-test with Welch’s correction. ***P < 0.001.

### DcpB is a c-di-GMP metabolizing protein

To understand the effect of DcpB on c-di-GMP metabolism, we measured the intracellular level of c-di-GMP of both wild-type C58 and Δ*dcpB* mutant strains carrying p*dcpB*, pGGAAF, pAAL, and pGGAAF-AAL expression plasmids. We also used a heterologous host, *E. coli* DH10B strain, to understand the enzymatic activity of each domain independent of endogenous *A. tumefaciens* regulatory control. In all three strain backgrounds, expression from plasmid-borne p*dcpB* reduced c-di-GMP levels (Fig. 4). Similarly, when expression was induced from pGGAAF, there was a reduction in c-di-GMP levels in all three backgrounds (Fig. 4), suggesting an active PDE domain. However, when the PDE domain is inactive and the GGDEF domain is intact, i.e., pAAL, there was a significant increase in c-di-GMP levels in the Δ*dcpB* mutant strain background (Fig. 4), indicating that in the absence of the PDE domain activity, the GGDEF domain is enzymatically active and capable of affecting c-di-GMP levels under certain conditions. A modest increase is apparent in the wild-type C58 background when the pAAL construct is induced, but does not reach the level of statistical significance. In the *E. coli* background, no difference in c-di-GMP levels is evident when pAAL is induced, suggesting *A. tumefaciens*-specific activity of the GGDEF domain. Expression from the pGGAAF-AAL construct showed no significant change in c-di-GMP in both wild-type C58 and Δ*dcpB* mutant strains (Fig. 4). Overall, our data suggests that DcpB is a c-di-GMP metabolizing protein with both DGC and PDE activity. However, under the experimental conditions tested PDE activity predominates, and DGC activity is masked or inhibited by the presence of an intact PDE domain.

**Fig 4.**
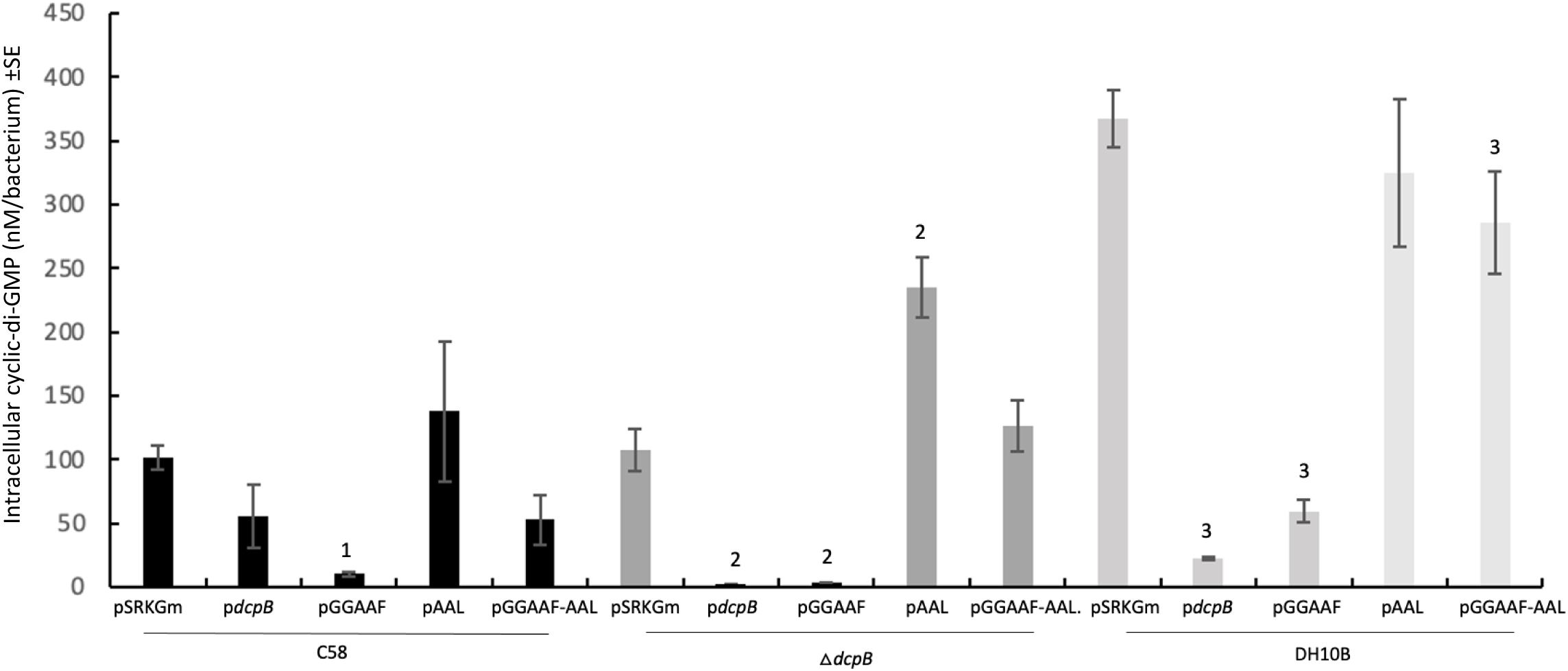
*dcpB* modulates intracellular levels of c-di-GMP. Intracellular levels of c-di-GMP were quantified using LC-MS/MS in either wild-type C58 *A. tumefaciens* (black bars) or the isogenic △*dcpB* strain (dark gray bars), as well as the heterologous *E. coli* DH10B host (light grey bars), carrying the same plasmid-borne alleles of *dcpB* from Figure 2. Statistical significance was determined using ANOVA Single Factor with Tukey Kramer. P < 0.05 for n = 3; ^1^compared to wild-type C58 pSRKGm, ^2^compared to △*dcpB* pSRKGm, ^3^compared to DH10B pSRKGm

### DcpB localizes to the poles and at mid-cell

During asymmetric cell division in *A. tumefaciens,* specific regulatory proteins differentially localize toward the older pole, younger pole, or at mid-cell (1, 17, 23). To study the localization of *dcpB*, we fused full-length DcpB to super-folder green fluorescent protein (DcpB-GFP) and performed epifluorescence microscopy on exponentially growing cells. DcpB-GFP expressed in both the wild-type and △*dcpB* strain backgrounds showed polar localization for 50% of cells compared to the total number of cells (Fig. 5, data not shown for △*dcpB* strain).

**Fig 5.**
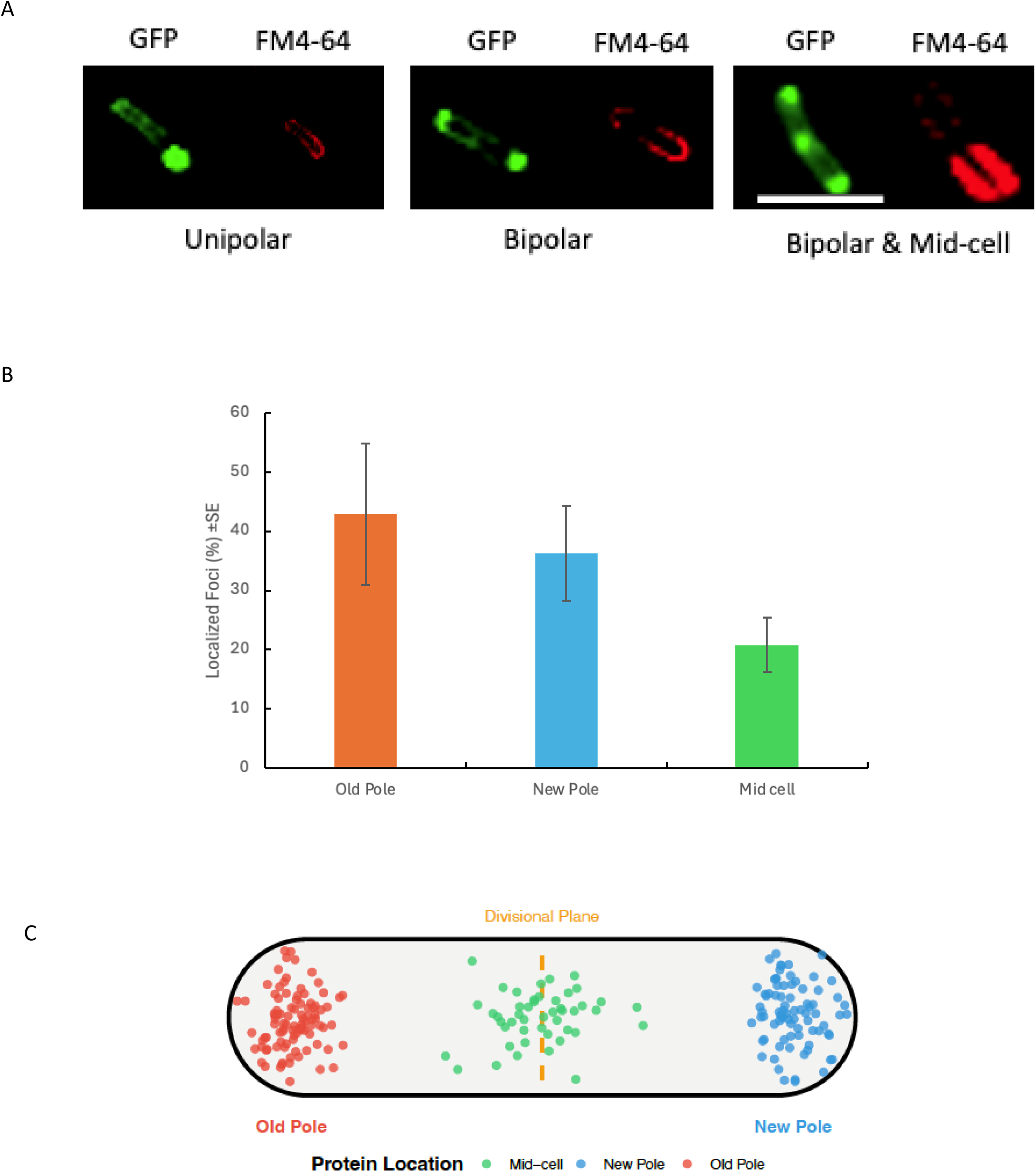
*dcpB*-GFP localizes at cell poles and mid-cell. **A.** Plasmid-borne expression of DcpB-GFP was induced with 250 µM IPTG in wild-type *A. tumefaciens* strain C58. Exponentially growing cultures were labelled with FM4-64 and imaged using phase contrast (not shown) and fluorescence microscopy. DcpB-GFP localization was compared with the FM4-64 labelling pattern to define GFP foci as unipolar, bipolar, or at mid-cell. Representative images are shown. Scale bar 2 µm. **B.** Quantification of DcpB-GFP focus localization for N = 226 foci, including foci in cells with multiple foci. 3 sets of data analyzed with 2-3 fields of view each. MicrobeJ plugin of ImageJ used to analyze the data. **C.** Schematic representation localization of the 226 foci quantified in panel B. Image made using R software.

Furthermore, to identify specific poles, we labeled the cell membrane with the intercalating lipophilic styryl fluorescent dye 4-64 (FM 4-64), as it has been demonstrated to robustly and preferentially label older poles in horseshoe patterns in *A. tumefaciens* (24–26). Our microscopy data included cell populations with unipolar, bipolar, and division plane (mid-cell) localization. Many cells had a combination of fluorescent foci resulting in unipolar plus division plane or bipolar and division plane localization (Fig. 5A). Out of the localized foci of DcpB-GFP, 43% localized in the old pole, 36% localized in the new pole, and 21% localized in the mid-cell in wild-type (Fig. 5B). On a population level, 50% of cells showed specific fluorescent foci indicating protein localization, while 50% showed only diffuse fluorescence; 36% showed unipolar, 5% showed bipolar, 5% showed divisional plane, 2% showed unipolar & divisional plane and 1% showed bipolar & divisional plane localization.

### DcpB expression is CtrA-dependent

Phosphorylated CtrA in *A. tumefaciens* interacts with the CtrA binding box (5’-TTAA-N_6_-GTTAAC-3’) present in the promoter region of genes encoding many cell cycle-regulated proteins, such as PdhS1 and CtrA, resulting variously in activation or repression of transcription (16, 17). The promoter region of DcpB (P*_dcpB_*) has two CtrA binding boxes at 301 and 147 bp upstream of the start codon with two CtrA binding sites (B.S.) each. The first CtrA binding box, 301 bp upstream, has the sequence 5’-TTAACACATTGTTAAC-3’, and the second CtrA binding box, 147 bp upstream, has the sequence 5’-TTTAGTGAAACTTAAT-3’, suggesting CtrA-dependent expression. We first decided to test the first binding box, as it has highly conserved CtrA binding site. Promoter activity of pRA301::P*_dcpB_* was evaluated in wild-type C58 and Δ*dcpB* mutant strain backgrounds via a beta-galactosidase assay. In addition, promoter activity of this construct was evaluated in both Δ*pdhS2* and Δ*divK* strain backgrounds, as these backgrounds are known to have altered CtrA activity in *A. tumefaciens* (17). Promoter activity of P*_dcpB_* was higher in the Δ*divK* strain and lower in the Δ*pdhS2* strain, compared to wild-type C58 (Fig. 6). P*_dcpB_* in the Δ*dcpB* mutant strain background showed similar promoter activity compared to wild-type C58 levels (Fig. 6). Our data are consistent with CtrA-dependent promoter activity of *dcpB* transcription.

**Fig 6.**
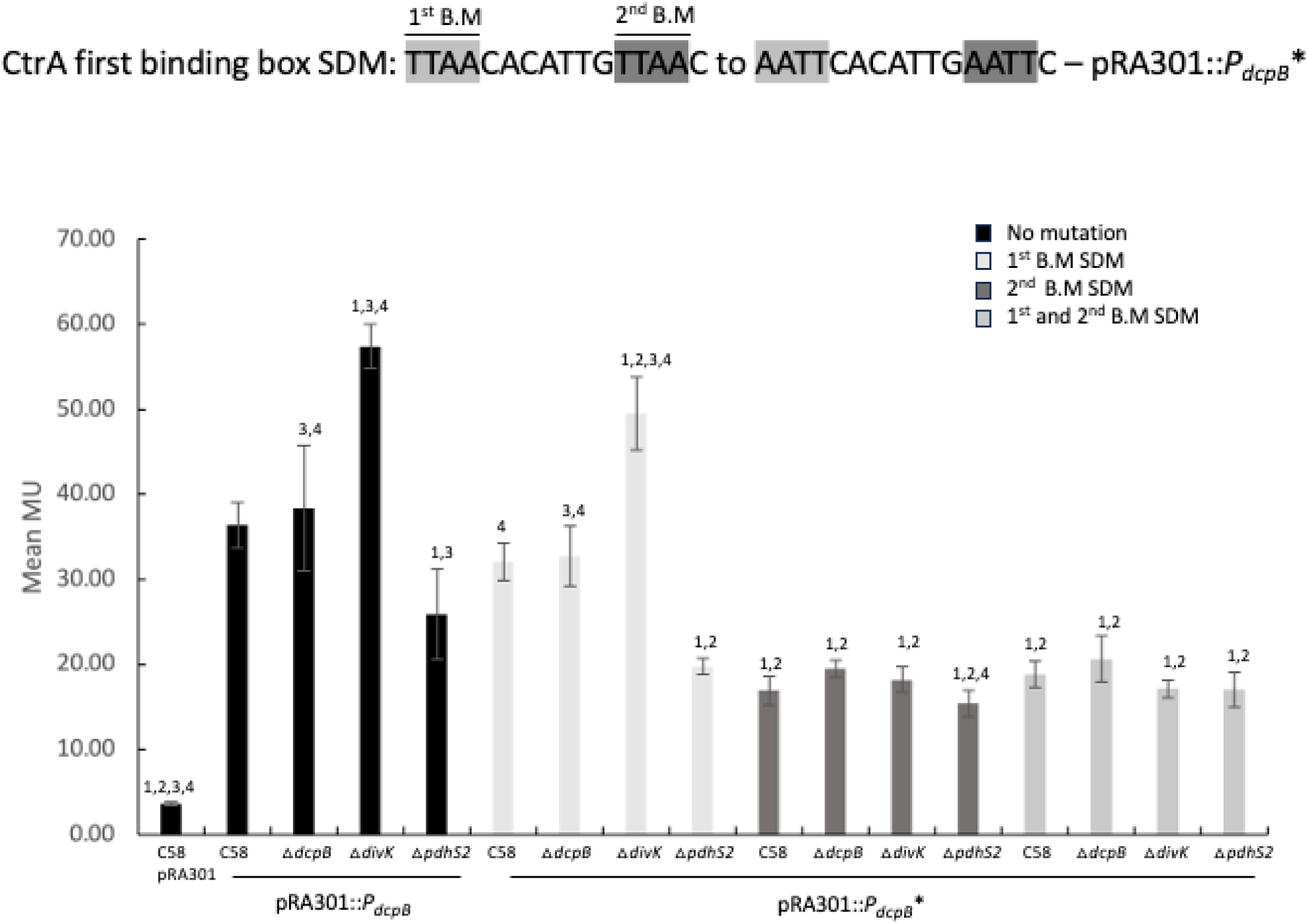
*dcpB* and *ctrA* expression is interdependent. **A.** Promoter activity of the wild-type *dcpB* promoter, or one of three mutated variants, was quantified as β-galactosidase activity from translational fusions of each promoter to *E. coli lacZ* in the pRA301 plasmid. Each construct was tested for activity in wild-type C58 *A. tumefaciens*, the △*dcpB* strain, and strains with increased (△*divK*) or decreased (△*pdhS2*) CtrA activity. The top image depicts the conserved CtrA binding site in the *dcpB* promoter, highlighting the two TTAA motifs. These motifs were changed via site-directed mutagenesis to AATT either singly or in combination, as indicated. *, reporter with 1^st^ (white bars), 2^nd^ (dark gray bars), or both (light gray bars) motifs altered. B.M, binding motifs. Statistical significance was determined using One Way ANOVA - Brown-Forsythe and Welch. P < 0.05 for n = 9; ^1^compared with wild-type C58 pRA301::*P_dcpB_*; ^2^compared with wild-type C58 pRA301::*P_dcpB*_* 1^st^ B.M. mutation; ^3^compared with wild-type C58 pRA301::*P_dcpB*_* 2^nd^ B.M. mutation; ^4^compared with wild-type C58 pRA301::*P_dcpB*_* dual B.M. mutation. MU, Miller units.

Based on the result of CtrA-dependent expression, we further wanted to test which CtrA potential binding motif within this CtrA binding box is required for cell cycle-dependent regulation of transcription from P*_dcpB_*. Using the pRA301::P*_dcpB_* construct, we mutated the CtrA first binding motif of the first binding site, 5’-TTAA-3’, to 5’-AATT-3’ and assayed promoter activity as before. Promoter activity decreased in the Δ*pdhS2* background and increased in the Δ*divK* background, similar to prior results with the wild-type promoter (Fig. 6). We further mutagenized the second CtrA binding motif, 5’- TTAA-3’ to 5’-AATT-3’, as well as simultaneously mutating both motifs, and assayed promoter activity. In both cases, there was a significant reduction in promoter activity compared to the wild-type promoter (Fig. 6). Moreover, cell cycle-dependent regulation of promoter activity was abrogated under these conditions. Based on these data, P*_dcpB_* has CtrA-dependent, and therefore cell cycle-dependent, expression, and the second binding motif within the first predicted CtrA binding box is required for this regulatory control.

### *thuR* genetically interacts with *dcpB*

To identify genetic interacting partners of *dcpB*, we performed transposon mutagenesis in the Δ*dcpB* strain and used suppressor screening to isolate mutants that were able to swim better than the Δ*dcpB* strain. A total of approximately 37,000 colonies were screened in three independent suppressor screens. Multiple independent hits were found in Atu3337 (*thuR*), a LacI family transcriptional regulator (Fig. 7A). We initiated characterization of *thuR* and its role in regulating developmental phenotypes. We started with a motility assay to verify the phenotype observed during screening for *thuR* transposon insertion in the Δ*dcpB* strain (Δ*dcpB* ThuR::Tn). Δ*dcpB* ThuR::Tn was motile compared to the Δ*dcpB* strain, confirming the suppressor screening phenotype (Fig. 7B). Furthermore, we wanted to know the phenotypic effect of Δ*dcpB* ThuR::Tn on attachment. We performed a static biofilm assay with the same strains as the motility assay. Compared to the mutant strain Δ*dcpB* and C58 wild-type strain, the transposon insertion strain had significantly increased adherent biomass (Fig. 7C).

**Fig 7.**
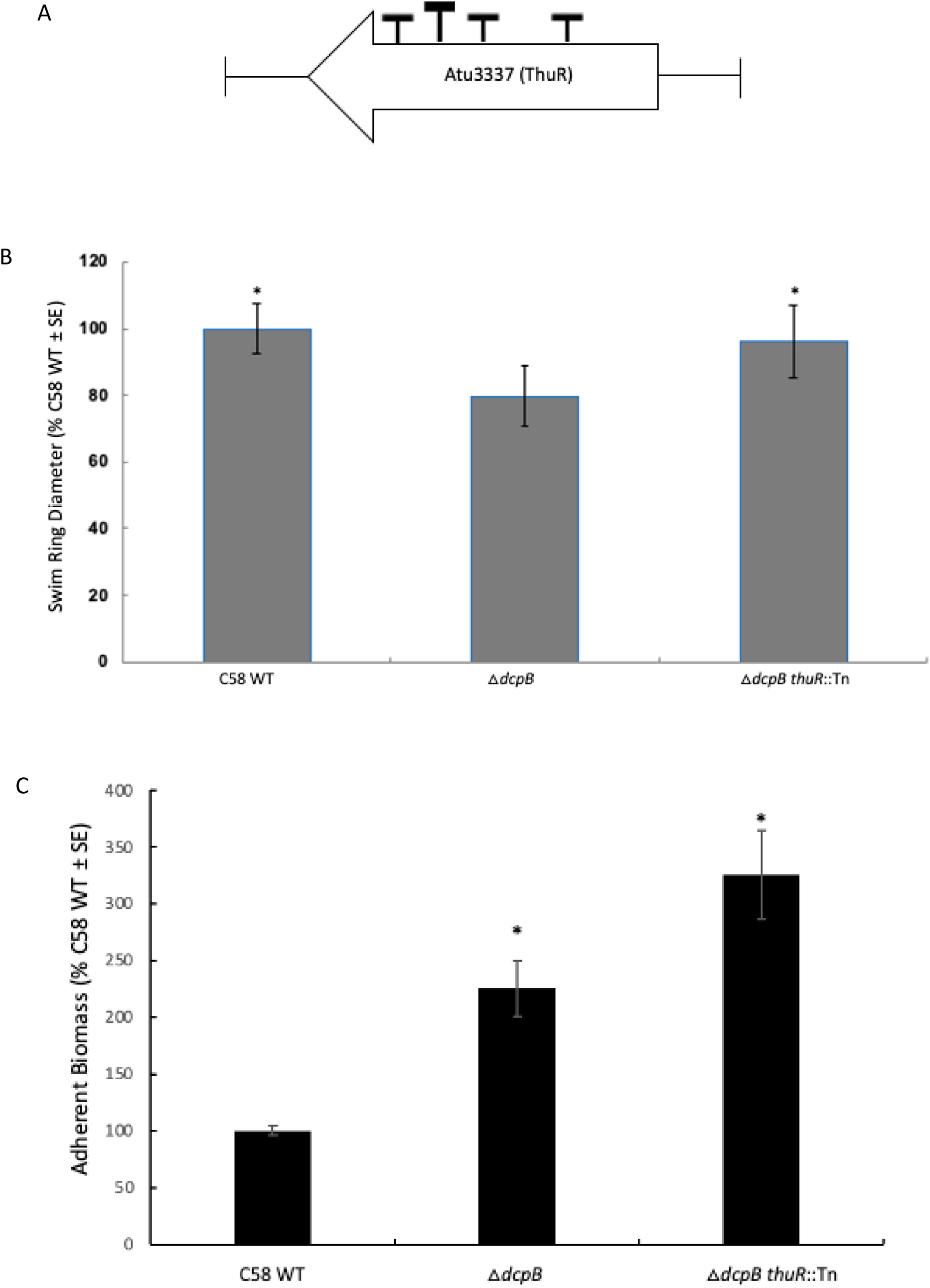
*thuR* inactivation suppresses the *dcpB* motility phenotype and exacerbates the *dcpB* biofilm phenotype. **A.** Transposon insertion sites in *thuR* (Atu3337) identified in △*dcpB*. Rectangles indicate independent insertion sites isolated from swim motility suppressor screening of approximately 37,000 colonies. **B.** Swimming motility was assessed as in Figures 1 and 2. Statistical significance was determined by Welch’s T-Test: Two-Tailed, Unequal Variance, compared to △*dcpB*. *P < 0.01 for n=9. **C.** Biofilm formation was assessed as in Figures 1 and 2. Statistical significance was determined by Student’s T-test comparing against wild-type C58. *P < 0.01 for n=9

## Discussion

Our data support the hypothesis that DcpB is primarily functioning as a PDE under our tested conditions. Overproduction of DcpB results in phenotypes consistent with reduced c-di-GMP levels, and absence of *dcpB* results in phenotypes consistent with increased c-di-GMP levels. Thus, in our experimental conditions, in *A. tumefaciens,* either DGC activity is inactive or PDE activity is dominant relative to DGC activity.

However, in the absence of PDE activity, DGC activity is unmasked. Another possibility may be that the DGC domain plays an intramolecular regulatory role. For example, the DGC domain may contribute to the activation of the PDE domain, similar to the BifA gene in *P. aeruginosa,* where the GGDEF functions as a GTP sensor, resulting in regulation of BifA EAL domain activity (27). In the well-studied model of *C. crescentus*, PdeA is a PDE with a degenerate GGDEF (GEDEF) domain lacking DGC activity that is still able to bind GTP (28, 29). PdeA is limited to the swarmer cell pole, where it limits the c-di-GMP level by antagonizing the activity of DGC DgcB and blocks cell differentiation (8). We propose that *A. tumefaciens* DcpB might be acting similarly to PdeA, as DcpB acts as a PDE by clearing the c-di-GMP level.

In the Δ*pdhS1*, Δ*pdhS2,* and Δ*visR* mutant strains, all of which are known to have reduced motility in the presence of wild-type levels of DcpB, no additional effect on motility was apparent when DcpB was overexpressed. However, biofilm formation was significantly decreased in all three strain backgrounds under these conditions. One reason for this disconnect could be due to artificial levels of DcpB leading to alterations in the global pool of c-di-GMP and that specific c-di-GMP effectors modulating biofilm formation are responsive to these changes. For motility, on the other hand, it may be local pools of c-di-GMP, synthesized and effective only at sites of synthesis, affect motility. Local c-di-GMP signaling involves spatially restricted changes in concentration of c-di-GMP that might affect cellular processes even though the global pool is not altered (30, 31). In *C. crescentus*, the DGC PleD is localized to the cell pole through its interaction with the polar organizing protein PodJ. PleD generates c-di-GMP locally at the pole, which then binds to nearby targets like PopA to control attachment, without significantly affecting the global c-di-GMP pool (8). It is telling that many studies using ectopic expression to study the effects of specific c-di-GMP-metabolizing proteins in *A. tumefaciens* rarely observe changes in motility during induction of either known DGC proteins or PDE proteins (12, 32) .

Sequence alignment of DcpB with well-studied DGCs and PDEs from other organisms highlights conservation of residues required for both DGC and PDE activity (Fig. S5). Furthermore, sequence alignment of the DGC catalytic motif of the 13 dual-domain proteins in *A. tumefaciens* revealed that there is no conserved I-site except in one protein, Atu4628; however, there is a conserved GTP and Mg^+2^ binding site in most of them required for the proper enzymatic activity of canonical DGC proteins (Fig. S6 & S7) (2, 33, 34).

The PdhS-DivK-CtrA regulatory pathway genes crucially affects *A. tumefaciens* cell cycle progression. The sensor kinase PdhS2 localizes to the younger pole after cytokinesis, and in the opposing pole DivJ and PdhS1 localize (17). In the case of DcpB, the unsynchronized cell population showed protein localization to the older pole, younger pole, and mid-cell (Fig. 5). These results suggest that the localization of DcpB is dynamic and depends on the particular stage of cell cycle. This is also the case for PdhS2 localization, where the protein localization changes as the cell divides to the newly generated young pole (17). Furthermore, downstream genes of *dcpB* include one of two predicted FtsK paralogs, Atu3210. *C. crescentus*, FtsK localizes at the mid-cell during cytokinesis resulting in new pole localization after completion of cell division (35). Having *ftsK* downstream of *dcpB* and *dcpB* mid-cell localization suggests the possibility that DcpB and FtsK both play a role in cell cycle progression in *A. tumefaciens*.

Based on our data, *dcpB* showed CtrA- and cell cycle-dependent expression (Fig. 6). Site-directed mutagenesis of the two potential binding motifs within the most upstream predicted CtrA binding site demonstrated that the first motif was not required while the second motif is required to support CtrA-dependent regulation from this promoter (Fig. 6). In the *dcpB* promoter, there are two predicted CtrA binding sites. In this work we have only evaluated the role played by the first, most upstream binding site. The second predicted CtrA binding site overlaps with the likely -35 region of the promoter, and most likely negatively regulates promoter activity; this will be tested in future work, but indicates complex regulation of *dcpB* promoter activity by CtrA.

Finally, we performed unbiased screening in the Δ*dcpB* strain and identified a LacI family repressor, ThuR, as genetically interacting with *dcpB* (Fig.7). Downstream of *thuR* lies the trehalose utilization operon, *thuEFGKAB* (Atu3338-Atu3343). In alphaproteobacteria, such as *S. meliloti* and *A. tumefaciens*, the *thu* operon (*thuEFGKAB*) is involved in production of an ABC transport system responsible for trehalose utilization, along with importing other sugars (36, 37). In *A. tumefaciens* and *S. meliloti* ABC transport systems are also regulated by the AbcR small regulatory RNAs (sRNA), AbcR1 and AbcR2, sibling sRNAs that target multiple components of ABC transport systems (38). RNA sequencing for a Δ*abcr1* mutant strain of *A. tumefaciens* C58 resulted in differential gene expression for *thuEFGKA* (38). How ThuR-dependent regulation of the *thu* operon intersects with DcpB activity is unclear. Uncovering a genetic interaction between *thuR* and *dcpB* provides a point of connection between the VtlR/AbcR and the PdhS-DivK-CtrA regulatory pathways, pointing to the potential centrality of ThuR as a potential virulence or fitness factor in *A. tumefaciens*.

## Materials and Methods

### Strains and Plasmids

All the strains, plasmids, and primers used in these studies are listed in Tables S1 - S3. AT minimal medium supplemented with 1% (w/v) glucose,15 mM ammonium sulfate, and 22 μM iron (II) sulfate (ATGN) was used for *A*. *tumefaciens* cultures, cultivated at 28 °C. *E. coli* was cultivated at 37 °C in lysogeny broth (LB). Antibiotics and the concentration used were as follows: (*E. coli*/*A. tumefaciens*), carbenicillin, 100 μg·mL^-1^/100 μg·mL^-1^; gentamicin, 25 μg·mL^-1^/150 μg·mL^- 1^; kanamycin, 25 μg·mL^-1^/150 μg·mL^-1^; and spectinomycin, 100 μg·mL^-1^/300 μg·mL^-1^. Isopropyl-β-D-thiogalactopyranoside (IPTG) 100 μM and 250 μM (microscopy) were used for protein induction.

### Strain and plasmid construction

Directional cloning was used for generating *dcpB* expression plasmids. Gibson assembly was used for generating DcpB::*sfGFP* in the pSRKKm expression plasmid. Site-directed mutagenesis (SDM) of *dcpB* was achieved using a modified QuikChange protocol (Agilent). For constructing the *lacZ* transcriptional reporter, the promoter region of *dcpB*, from 500 bp upstream of the start codon to 9 bp downstream, was cloned into pRA301, which includes a promoterless *E. coli lacZ* gene.

### Non-polar deletions

The deletion protocol was performed as published previously (39). Briefly, splicing by overlap extension (SOE) PCR was used to amplify approximately 500 bp upstream of the *dcpB* locus and approximately 500 bp downstream. Upstream and downstream amplicons were then used as both template and primer to splice the two together. Amplicons were sub-cloned into pNPTS138. The SOE constructs were delivered into *A. tumefaciens* strain C58 by conjugation. Transconjugants were selected by plating on ATGN plates with 300 μg·mL^-1^ kanamycin. Transconjugants were counterselected by plating onto AT plates supplemented with 5% sucrose. Counterselected recombinants were patched onto ATGN plates with 300 μg·mL^-1^ kanamycin and AT plates supplemented with 5% sucrose. Km^S^ Suc^R^ candidates were streak purified and verified by PCR.

### Static biofilm assay

Overnight cultures were set up at 28 °C in ATGN media along with appropriate antibiotics, and 100 μM IPTG if needed. The following morning, cultures were sub-cultured to an OD_600nm_ of 0.1 and incubated at 28 °C until the culture reached 0.25 – 0.8 OD_600nm_. Cultures were then diluted down to an OD_600nm_ of 0.05. Static biofilm assay was conducted in 12-well plates with 3 mL of inoculum. To analyze the biofilm formation, each coverslip was removed from the well, rinsed with deionized water, and immersed into 0.1% (w/v) crystal violet solution. Rinsed coverslips were then placed in 1 mL of 33% acetic acid to solubilize the crystal violet stain.

### Motility assay

0.25 – 0.3% (w/v) loose agar plates were prepared with ATGN supplemented with appropriate antibiotics and IPTG. A single colony of the test strain was inoculated in the center of the plate by using sterile wooden sticks. Plates were incubated at room temperature for seven days.

### Growth curve

5 mL overnight cultures were inoculated in ATGN media and incubated at 28 °C. The next morning, all the strains were sub-cultured to an OD_600nm_ of 0.05. 100 μM IPTG was added when appropriate. Cultures were incubated with shaking at 28 °C, and OD_600nm_ readings were taken every 2 hours for 12 hours, and then again at 24 and 48 hours.

### Potato tumor assay

To study tumor formation, test strains were cultured in 5 mL of ATGN overnight with shaking at 28 °C. The following morning, OD_600nm_ was measured, and each culture was sub-cultured to an OD_600nm_ of 0.01. All the cultures were then incubated with shaking at 28 °C to mid-exponential phase. Cultures were diluted down to an OD_600nm_ of 0.05. Sterile potato discs were placed on 1.5% agar plates with no additional nutrients. Each disc was inoculated with 10 μL of the test strain. The plates were sealed with parafilm and stored at room temperature for 4 weeks.

### Transposon mutagenesis and motility suppressor isolation

Transposon mutagenesis was achieved using plasmid pFD1, carrying the *mariner* family transposon *Himar1*, mobilized via conjugation by donor *E. coli* S17-1/λ*pir* and recipient strains *A. tumefaciens* Δ*dcpB* or *A. tumefaciens* Δ*pdhS1*. Strains bearing transposon insertions that suppressed the motility phenotype of the parent strain were isolated as follows. 10 μL of the mutant library was inoculated into the center of swim agar plates (0.25% w/v) and incubated at room temperature. Samples were collected from the flares or edges of the resulting swim rings and streak purified onto ATGN 300 μg·mL^-1^ kanamycin to isolate single colonies. Touchdown PCR was performed to identify the site of *Himar1* insertion.

### Beta-galactosidase (β-gal) assay

A modified β-galactosidase assay was performed to measure the promoter activity of derivatives of the *dcpB* or *ctrA* promoters as previously published (40). Test strains carrying transcriptional reporter constructs were grown overnight in ATGN media with appropriate antibiotics at 28 °C and sub-cultured to an OD_600nm_ of 0.1 the following morning. The cultures were incubated at 28 °C with shaking until mid-exponential phase, 0.3 – 0.6 OD_600nm_. In a sterile microfuge tube, 0.5 mL of the culture and 0.5 mL of Z-buffer (60mM Na^2^HPO^4^·7H_2_0, 40mM NaH^2^PO_4_, 10mM KCl, 1mM MgSO^4^·7H_2_O, pH 7.0) were mixed. Two drops of 0.05% sodium dodecyl sulfate and three drops of chloroform were added, and the mixture was vortexed. 0.1 mL of 4 mg·mL^-1^ of *O*-nitrophenyl-*β*-D-galactopyranoside (ONPG) made in Z-buffer was added, and a timer was started. Once the solutions started turning light yellow, the reactions were stopped by adding 1 M sodium carbonate, and the time (*t*) was recorded. The absorbance at 420 nm was measured in 96-well plates for each reaction.

### Microscopy

For DcpB-GFP localization, overnight cultures were grown at 28 °C with shaking in ATGN media with 150 μg·mL^-1^ kanamycin and 250 μM IPTG. Cultures were sub-cultured to an OD_600_ of 0.1 and allowed to grow until an OD_600_ of 0.4 – 0.6. 8 μg·mL^-1^ FM 4-64 was added in 1 ml of culture and stained for 8 mins at RT in the dark to label the cell membrane (24, 41). The labelled cell pellet was collected by centrifugation at 7,000 x g for 5 mins and resuspended in 1 ml of ATGN. The cells were washed a total of four times. 1 μL was pipetted onto a 1% agarose pad made in ATGN media. Images were acquired using a K3M digital camera attached to a DMi8 inverted microscope, an EL6000 mercury halide external light source, phase contrast optics, and epifluorescence, with an HC PL Fluotar 100x/1.32 oil immersion Ph3 objective and GFP filter (Leica Microsystems). The images collected were analyzed using MicrobeJ (42), an ImageJ plugin.

### c-di-GMP Measurement

*E. coli* DH10B strains were grown in LB media with 25 μg·mL^-1^ gentamicin plus 100 μM IPTG and incubated at 37 °C with shaking to an OD_600_ above 0.5. *A. tumefaciens* strains were grown in ATGN media with 150 μg·mL^-1^ gentamicin plus 100 μM IPTG to stationary phase, OD_600_ > 1.0. Culture density was determined for all the samples, and the volumes of the samples were adjusted to make sure each sample had a similar number of cells. Cells were collected by centrifugation at 12,500 x *g* for 5 min at 4 °C. The pellets were resuspended in 100 μL and 250 μL of extraction buffer (0.1 M formic acid in a 40:40:20 ratio of acetonitrile:methanol:water), chilled at -20 °C, for *E. coli* and *A. tumefaciens,* respectively (12, 43, 44). The extractions were incubated at -20 °C for 20 mins and centrifuged for 15 mins at 15,000 x *g* at 4 °C. The supernatants were transferred into a new tube and dried. Dried samples were redissolved in 100 μL of mobile solvent phase A. LC-MS/MS was used to analyze 10 μL of each sample on a Quattro Premier XE mass spectrometer (Waters Corporation) coupled with an Acquity Ultra Performance LC system (Waters Corporation) (12, 40). c-di-GMP standards (Axxora) were dissolved in the extraction buffer to create a standard curve.

## Acknowledgments

This work was funded by NSF CAREER Award 2238568 and NIFA CAREER Award 2023-67014-40246 (JEH), and NIH Awards GM139537 and AI158433 (CMW). The authors thank Zachary Hassler for assisting with suppressor screening.

---

**Table S1:**
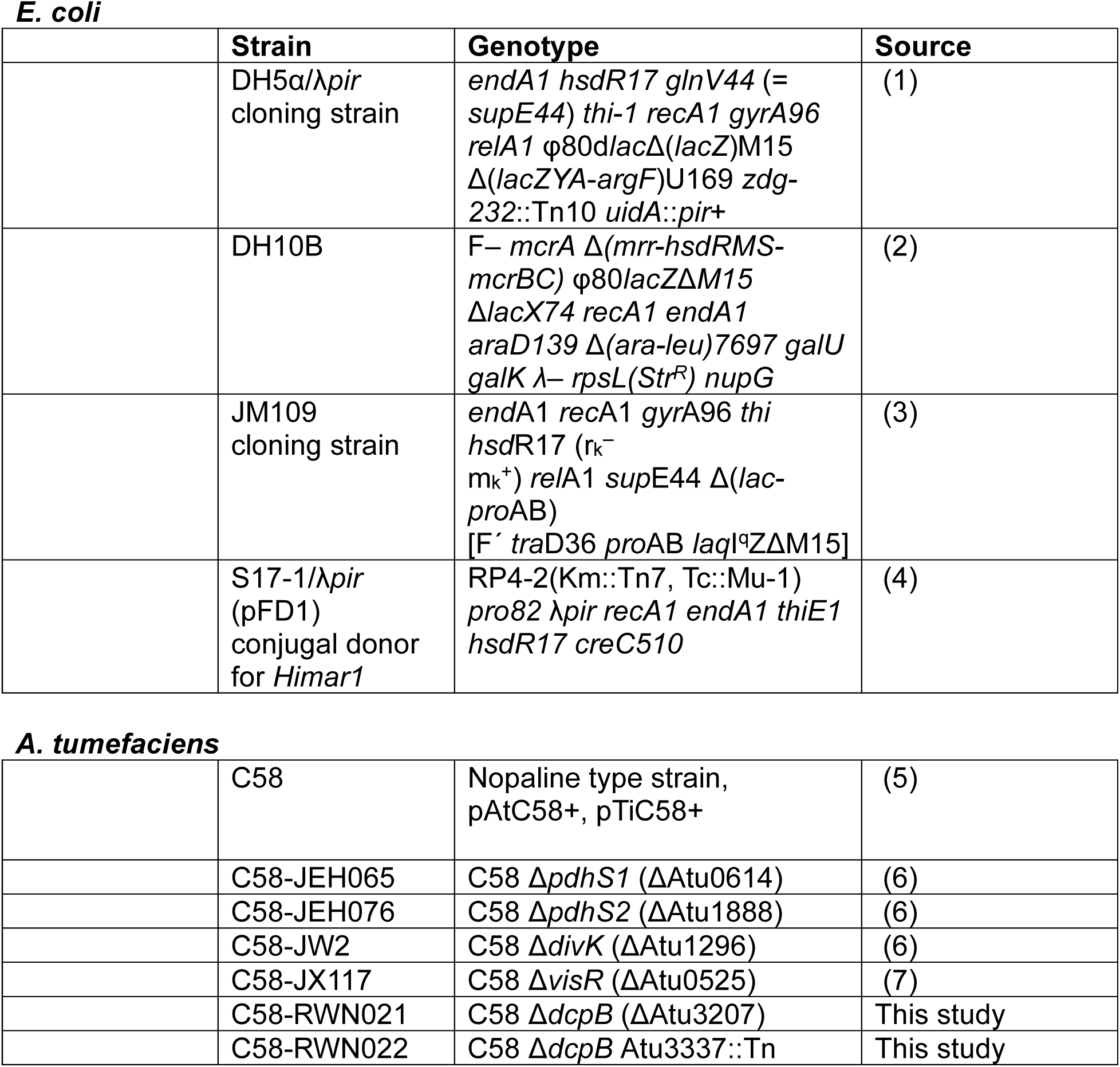
List of strains used in this study.

|  | Strain | Genotype | Source |
| --- | --- | --- | --- |
| | DH5α/*λpir*<br>cloning strain | *endA1 hsdR17 glnV44* (= *supE44*) *thi-1 recA1 gyrA96 relA1* $\phi 80\text{dlac}\Delta(\text{lacZ})\text{M15}$ $\Delta(\text{lacZYA-argF})\text{U169 zdg-232::Tn10 uidA::pir+}$ | (1) |
| | DH10B | F<sup>−</sup> *mcrA* $\Delta(\text{mrr-hsdRMS-mcrBC})$ $\phi 80\text{lacZ}\Delta\text{M15}$ $\Delta\text{lacX74 recA1 endA1 araD139}$ $\Delta(\text{ara-leu})7697$ *galU galK* $\lambda$− *rpsL(Str<sup>R</sup>) nupG* | (2) |
| | JM109<br>cloning strain | *endA1 recA1 gyrA96 thi hsdR17* ($r_k^- m_k^+$) *relA1 supE44* $\Delta(\text{lac-proAB})$ [F' *traD36 proAB laqI<sup>q</sup>Z*$\Delta\text{M15}$] | (3) |
|  | S17-1/*λpir*<br>(pFD1)<br>conjugal donor<br>for *Himar1* | RP4-2(Km::Tn7, Tc::Mu-1) *pro82 λpir recA1 endA1 thiE1 hsdR17 creC510* | (4) |
| ***A. tumefaciens*** |  |  |  |
|  | C58 | Nopaline type strain, pAtC58+, pTiC58+ | (5) |
| | C58-JEH065 | C58 $\Delta\text{pdhS1}$ ($\Delta\text{Atu0614}$) | (6) |
| | C58-JEH076 | C58 $\Delta\text{pdhS2}$ ($\Delta\text{Atu1888}$) | (6) |
| | C58-JW2 | C58 $\Delta\text{divK}$ ($\Delta\text{Atu1296}$) | (6) |
| | C58-JX117 | C58 $\Delta\text{visR}$ ($\Delta\text{Atu0525}$) | (7) |
| | C58-RWN021 | C58 $\Delta\text{dcpB}$ ($\Delta\text{Atu3207}$) | This study |
| | C58-RWN022 | C58 $\Delta\text{dcpB}$ *Atu3337::Tn* | This study |

**Table S2:** List of plasmids used in this study.

| Plasmid name | Characteristics | Source |
| --- | --- | --- |
| pGEM-T Easy | PCR cloning vector; Amp<sup>R</sup> | Promega |
| pNPTS138 | ColE1 origin; *sacB*; Km<sup>R</sup> | Gift of M. Alley |
| pRA301 | Broad host range, carrying promoterless *lacZ*; Spec<sup>R</sup> | (8) |
| pSRKGm | Broad host range *P<sub>lac</sub>* expression vector; *lacI*Q; Gm<sup>R</sup> | (9) |
| pSRKKm | Broad host range *P<sub>lac</sub>* expression vector; *lacI*Q; Km<sup>R</sup> | (9) |
| pJEH122 | pRA301 carrying *P<sub>ctrA</sub>-lacZ* | (10) |
| pJEH166 | pSRKGm carrying *P<sub>lac</sub>-dcpB*<sup>E311A E312A</sup> (pGGAAF) | This study |
| pJEH167 | pSRKGm carrying *P<sub>lac</sub>-dcpB*<sup>E441A</sup> (pAAL) | This study |
| pJEH177 | pGEM-T Easy carrying full-length *dcpB* | This study |
| pJEH191 | pSRKGm carrying *P<sub>lac</sub>-dcpB* | This study |
| pJEH203 | pGEM-T Easy carrying SOE construct for *dcpB* deletion | This study |
| pJEH204 | pNPTS138 carrying SOE construct for *dcpB* deletion | This study |
| pJEH225 | pSRKKm carrying *P<sub>lac</sub>-dcpB::sfGFP* | This study |
| pRWN018 | pRA301 carrying *P<sub>dcpB</sub>*<sup>TTAA-301-298AATT</sup>-*lacZ* (SDM of 1<sup>st</sup> CtrA half-site) | This study |
| pRWN020 | pSRKGm carrying *P<sub>lac</sub>-dcpB*<sup>E311A E312A E441A</sup> (pGGAAF-AAL) | This study |
| pRWN028 | pRA301 carrying *P<sub>dcpB</sub>-lacZ* | This study |
| pRWN035 | pRA301 carrying *P<sub>dcpB</sub>*<sup>TTAA-301-298AATT TTAA-290-287AATT</sup>-*lacZ* (SDM of both CtrA half-sites) | This study |
| pRWN037 | pRA301 carrying *P<sub>dcpB</sub>*<sup>TTAA-290-287AATT</sup>-*lacZ* (SDM of 2<sup>nd</sup> CtrA half-site) | This study |

**Table S3:**
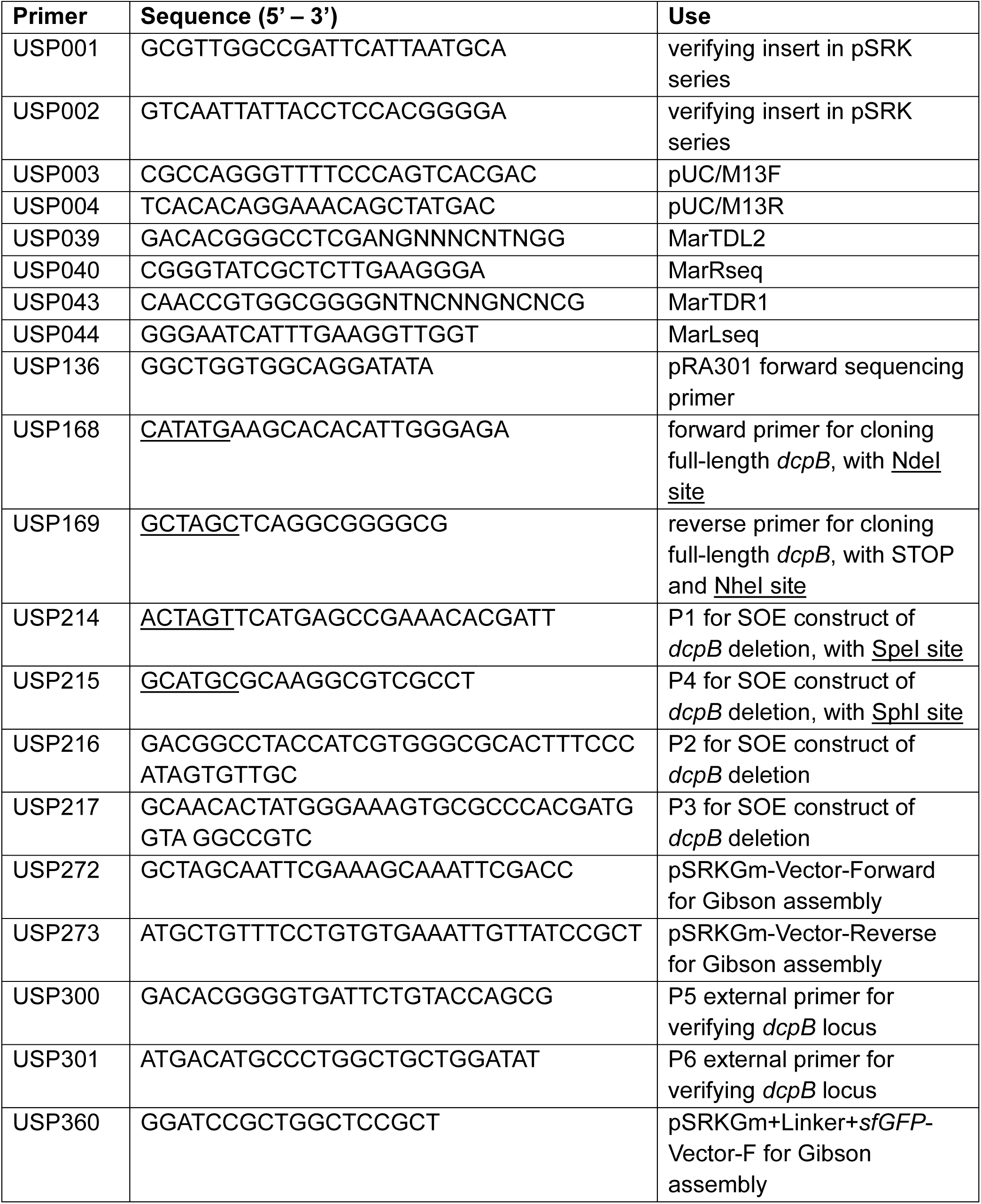
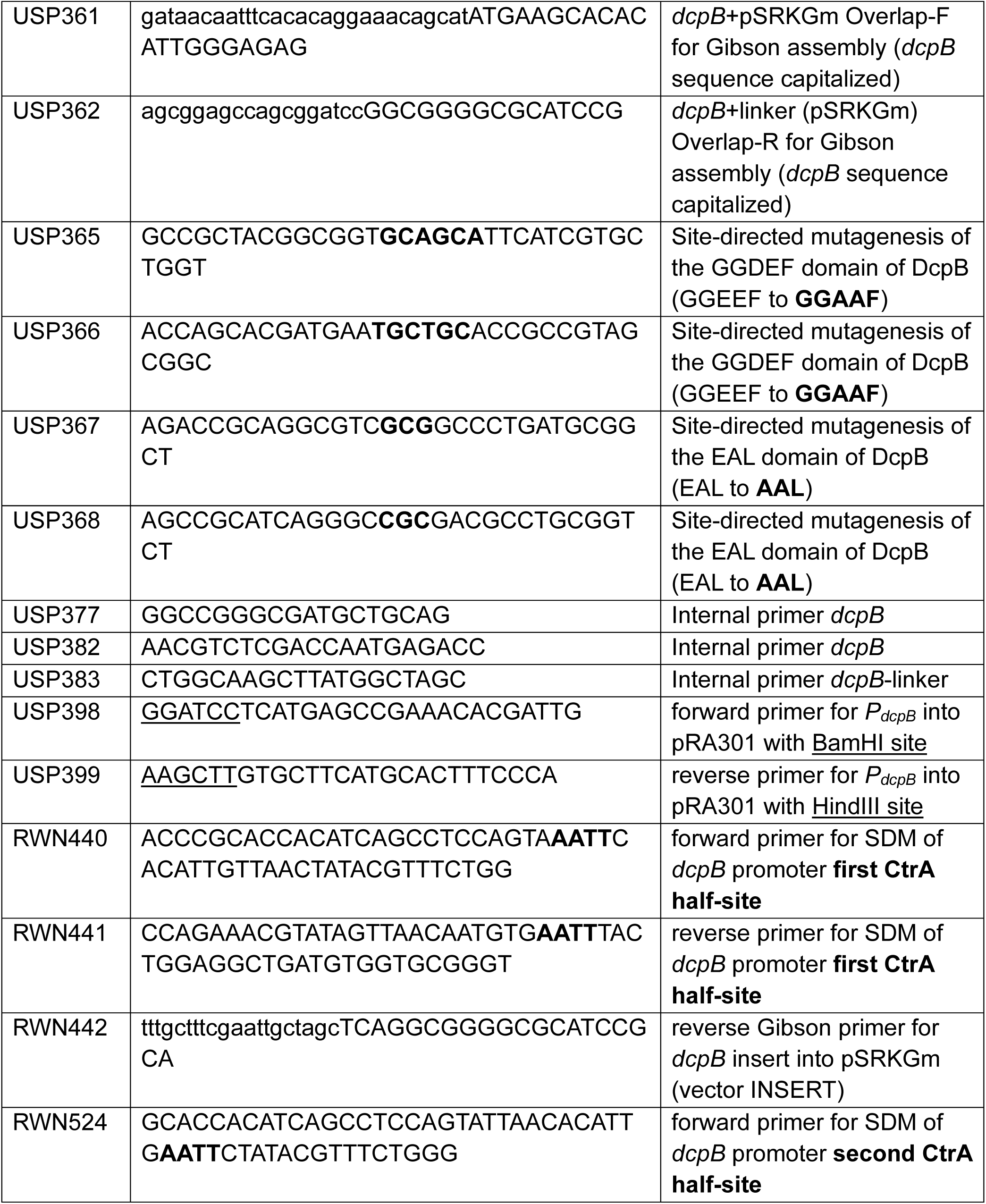
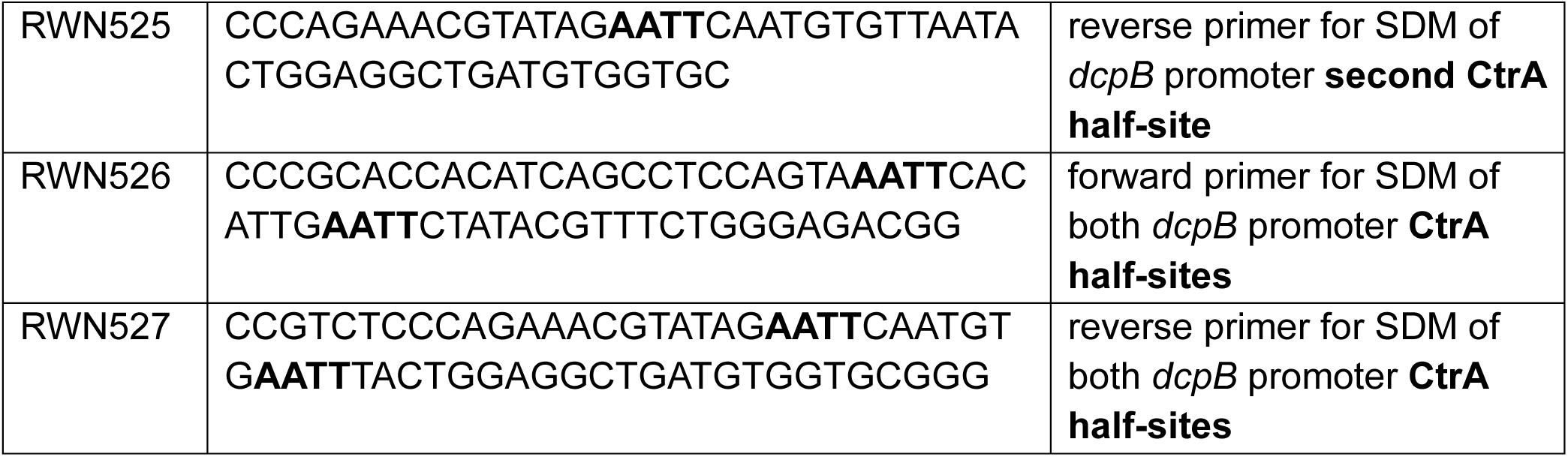
List of DNA primers used in this study.

**Fig S1.**
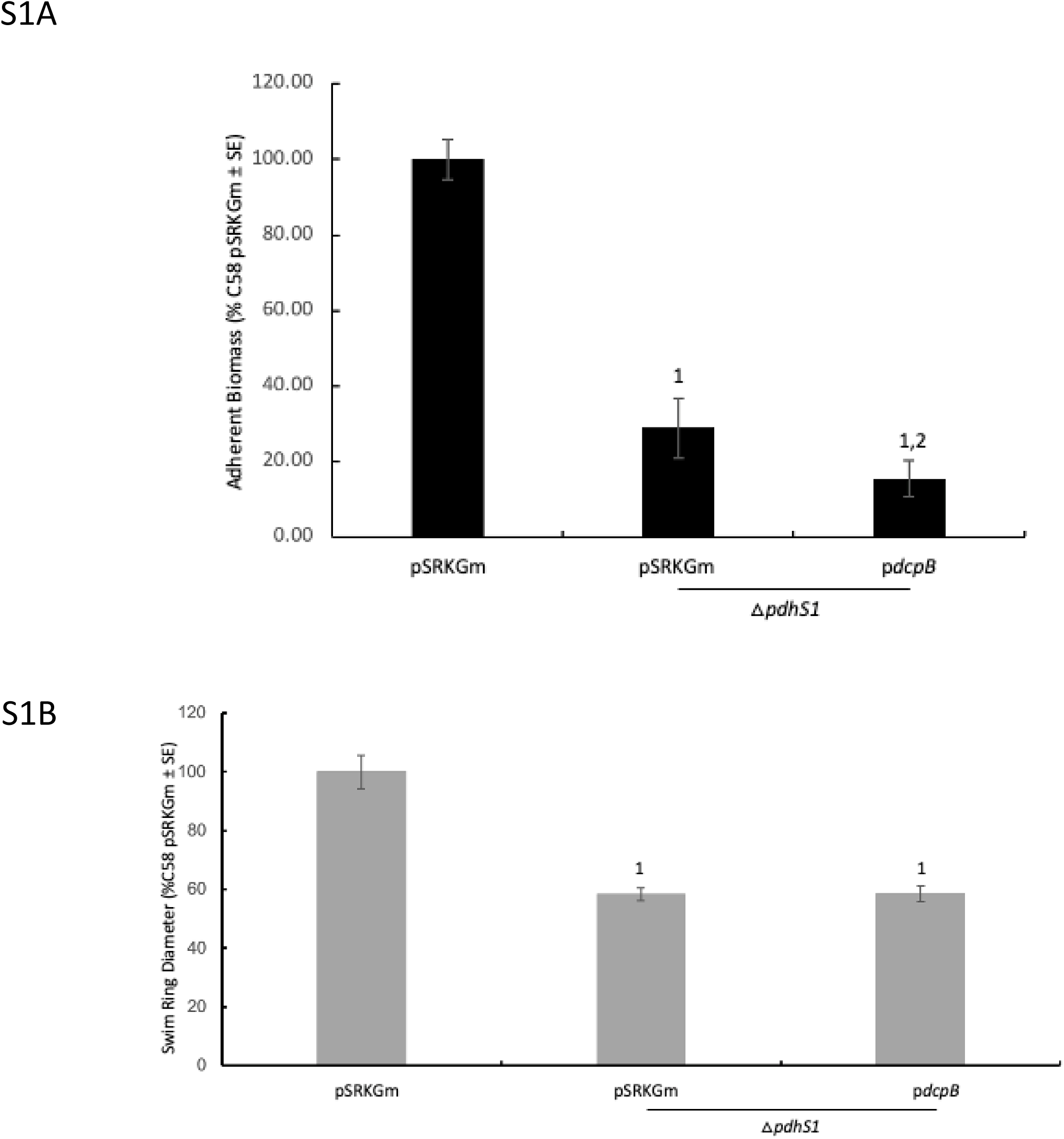
*dcpB* expression reduces biofilm formation but has no effect on motility in the △*pdhS1* strain background. **A.** Biofilm formation was assessed as in Figures 1 and 2. Statistical significance was determined using t-test: two-sample assuming equal variances. ^1^P < 0.01 compared to wild-type C58 pSRKGm and ^2^compared to △*pdhS1* pSRKGm. N=9. **B.** Swimming motility was assessed as in Figures 1 and 2. Day seven data are shown. All strains were induced with 100 µM IPTG. Statistical significance was determined using T-test: two-tailed, unequal variances. ^1^P < 0.01 compared to wild-type C58 pSRKGm. N=9.

**Fig S2.**
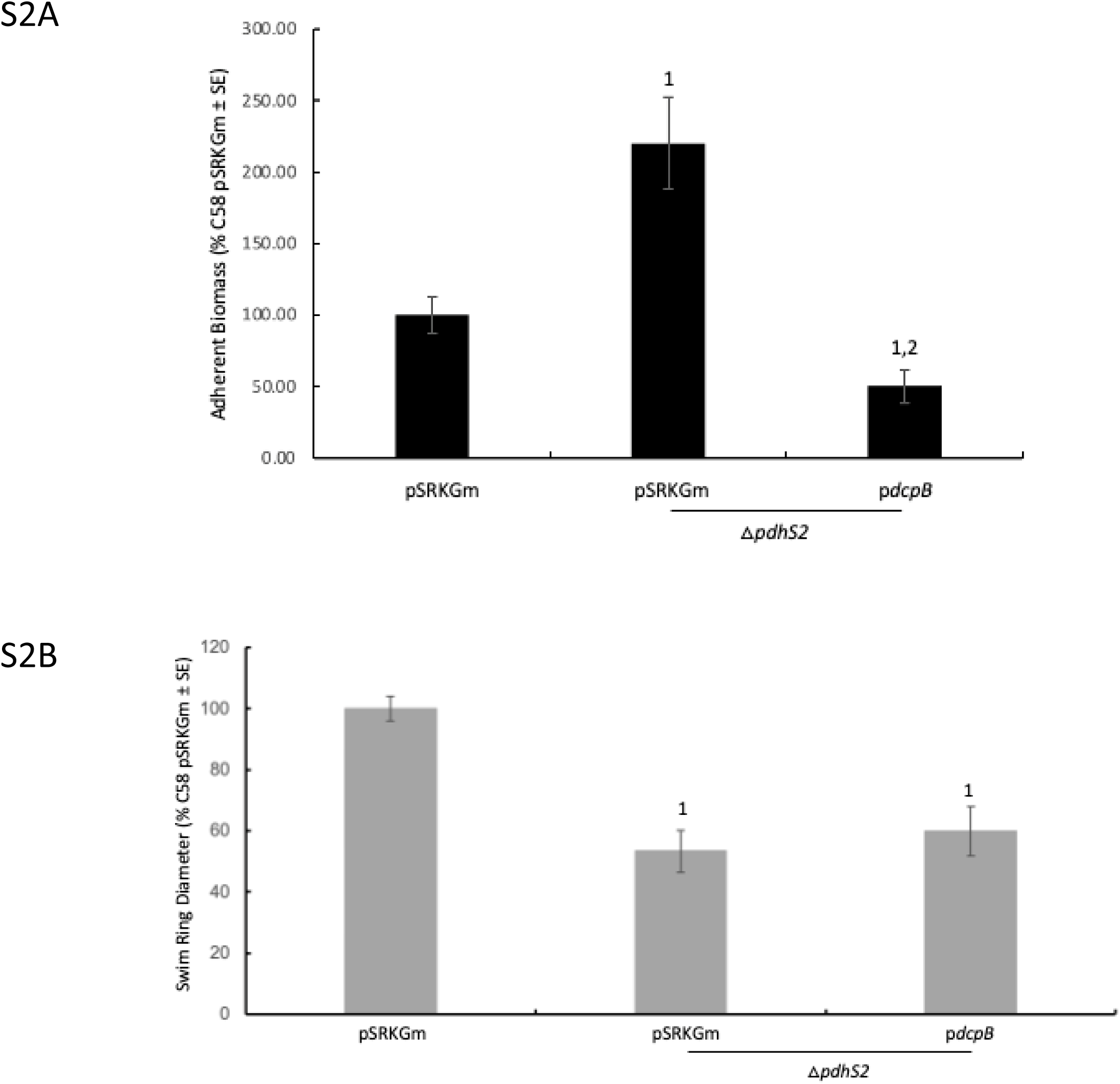
*dcpB* expression reduces biofilm formation but has no effect on motility in the △*pdhS2* strain background. **A.** Biofilm formation was assessed as in Figures 1 and 2. Statistical significance was determined using t-test: two-sample assuming equal variances. ^1^P < 0.01 compared to wild-type C58 pSRKGm and ^2^compared to △*pdhS2* pSRKGm. N=9. **B.** Swimming motility was assessed as in Figures 1 and 2. Day seven data are shown. All strains were induced with 100 µM IPTG. Statistical significance was determined using T-test: two-tailed, unequal variances. ^1^P < 0.01 compared to wild-type C58 pSRKGm. N=9.

**Fig S3.**
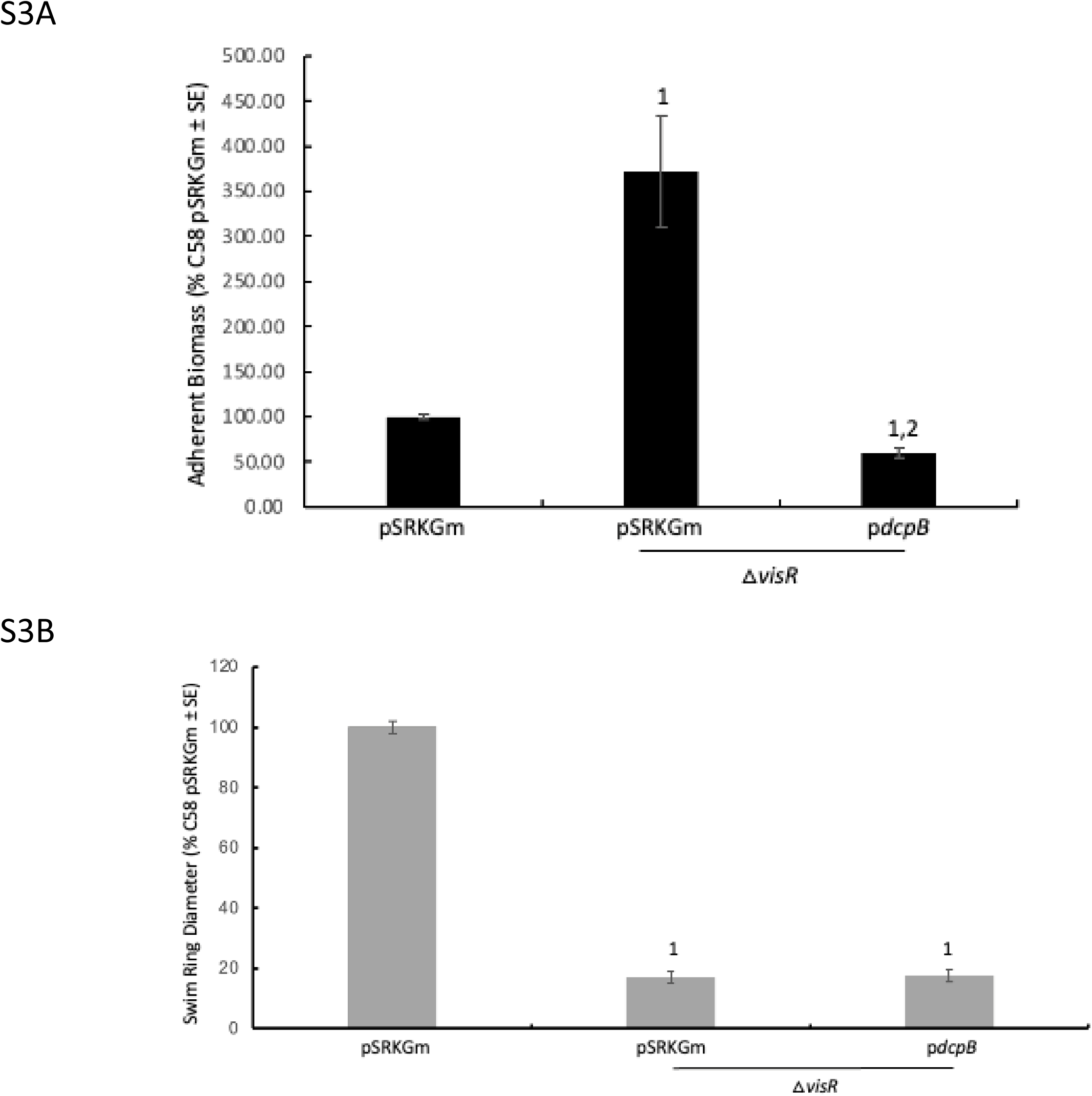
*dcpB* expression reduces biofilm formation but has no effect on motility in the △*visR* strain background. **A.** Biofilm formation was assessed as in Figures 1 and 2. Statistical significance was determined using one-way ANOVA with Brown-Forythe and Welch test. ^1^P < 0.05 compared to wild-type C58 pSRKGm and ^2^compared to △*visR* pSRKGm. N=6. **B.** Swimming motility was assessed as in Figures 1 and 2. Day seven data are shown. All strains were induced with 100 µM IPTG. Statistical significance was determined using T-test: two-tailed, unequal variances. ^1^P < 0.01 compared to wild-type C58 pSRKGm. N=6.

**Fig. S4.**
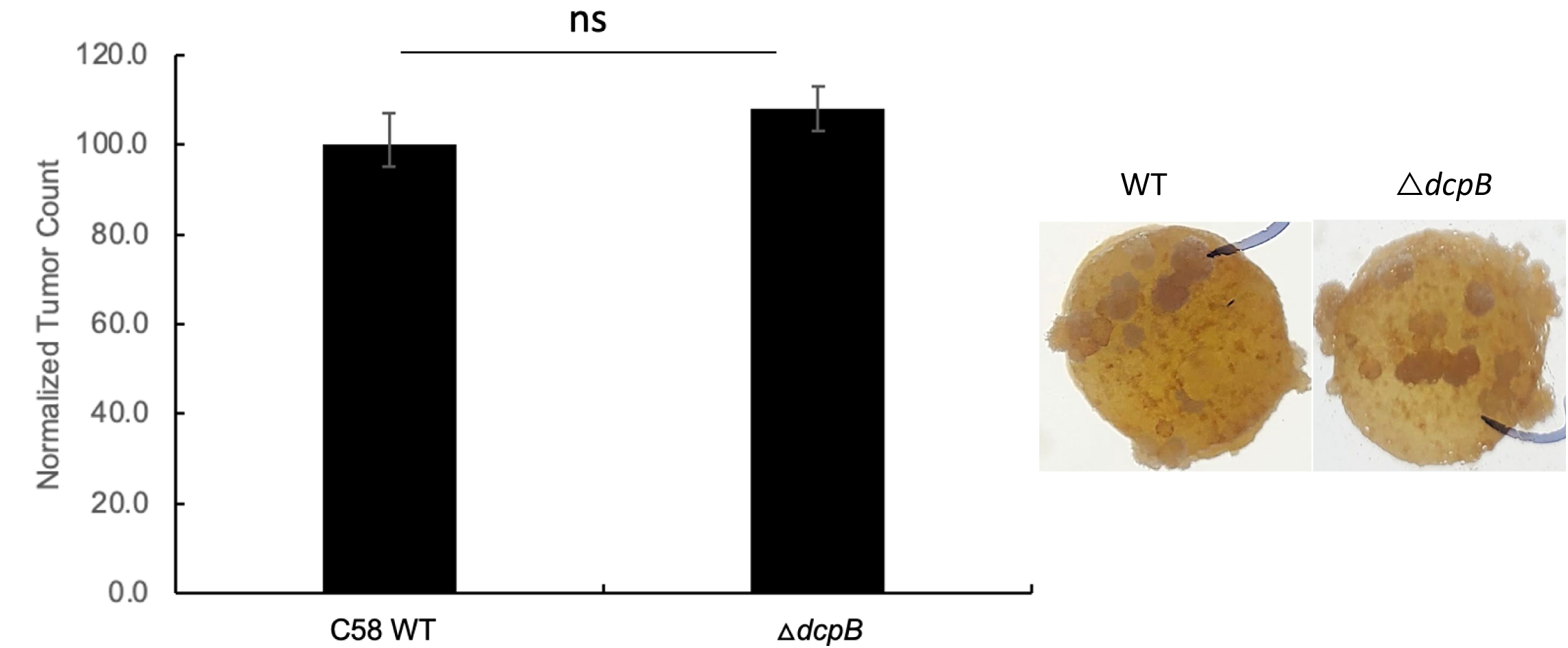
Absence of *dcpB* results in minimal affect on tumor formation. Potato disc tumor assay was performed to measure the tumor formation. Tumor count was determined weekly for four weeks. Data shown are for week four. There was no significant difference based on Paired t-test in tumor formation for week four tumors. Representative images are included of potato discs with tumors resulting from infection with wild-type C58 and the △*dcpB* strain. ns, not significant.

**Fig. S5.**
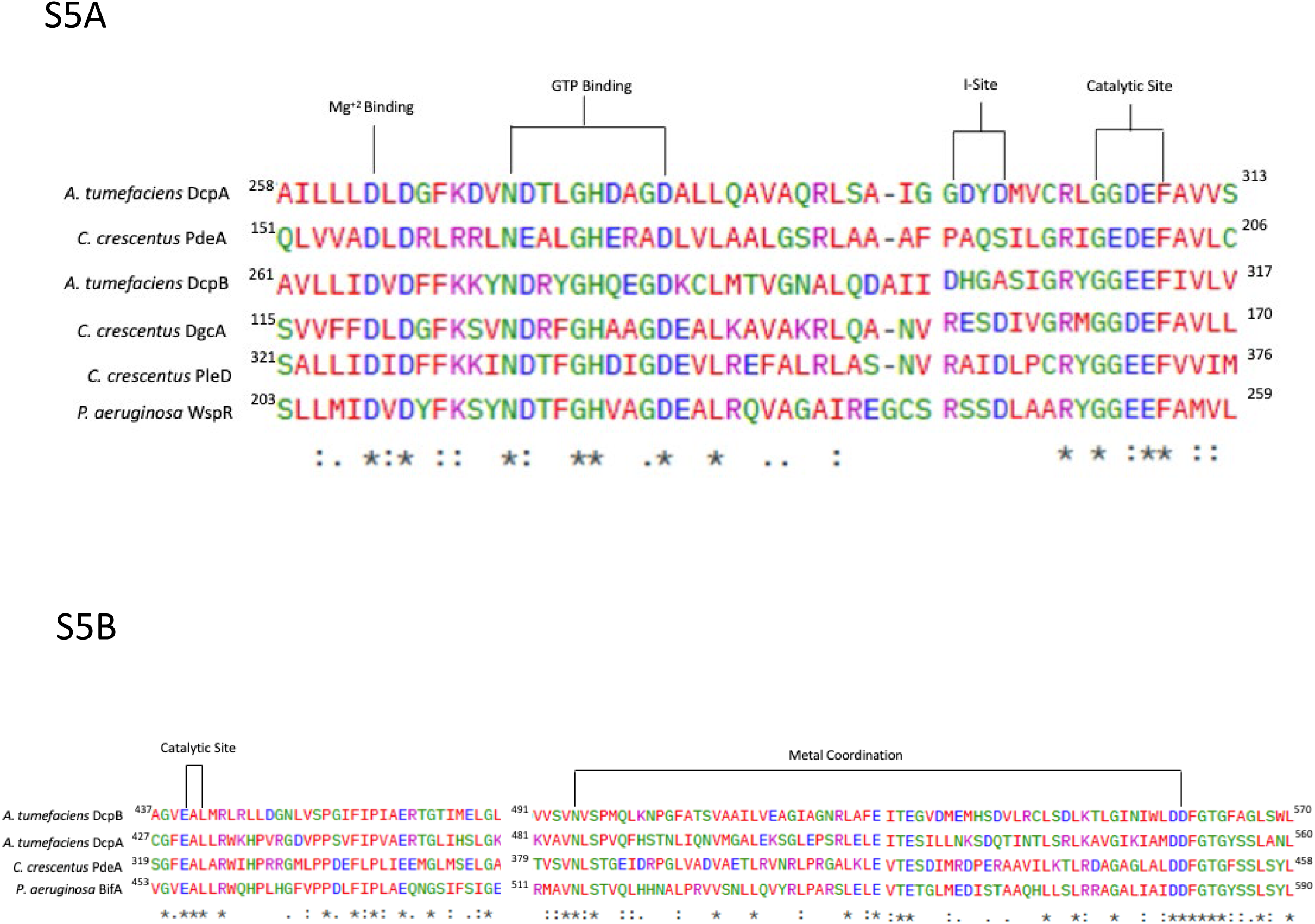
DcpB maintains important conserved active site features for both GGDEF and EAL domains. **A.** Sequence alignment for the diguanylate cyclase domain showing Mg^+2^ binding site, GTP binding, allosteric inhibitory I site, and GGDEF catalytic site among representative diguanylate cyclase proteins from *A. tumefaciens*, *C. crescentus*, and *P. aeruginosa*. **B.** Sequence alignment for the phosphodiesterase domain showing EAL catalytic site along with other conserved metal binding residue in *A. tumefaciens, C. crescentus*, and *P. aeruginosa* . Alignments performed using Clustal Omega EMBL-EBI.

**Fig. S6.**
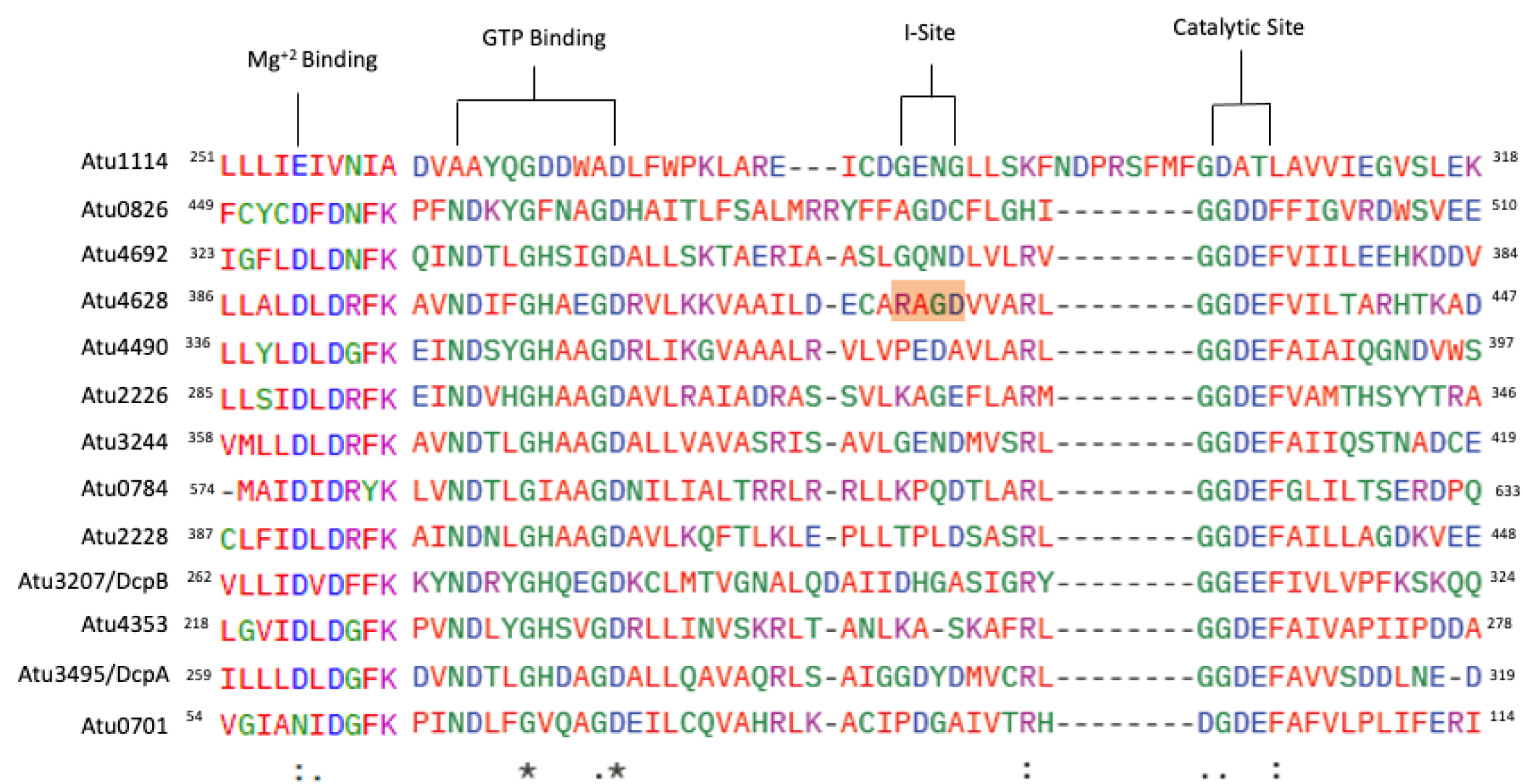
Dual-domain diguanylate cyclase/phosphodiesterase proteins from *A. tumefaciens* show loss of conservation of the inhibitory I site. Sequence alignment of the 13 predicted dual-domain proteins highlighting conserved elements of the diguanylate cyclase domain. The Mg^+2^ binding, GTP binding, and catalytic sites are shown in the alignment. The inhibitory I-site (RxxD) was only observed in Atu4692, orange color. Atu1114 was used as the reference sequence Protein sequences were aligned with Clustal Omega EMBL-EBI.

**Fig. S7.**
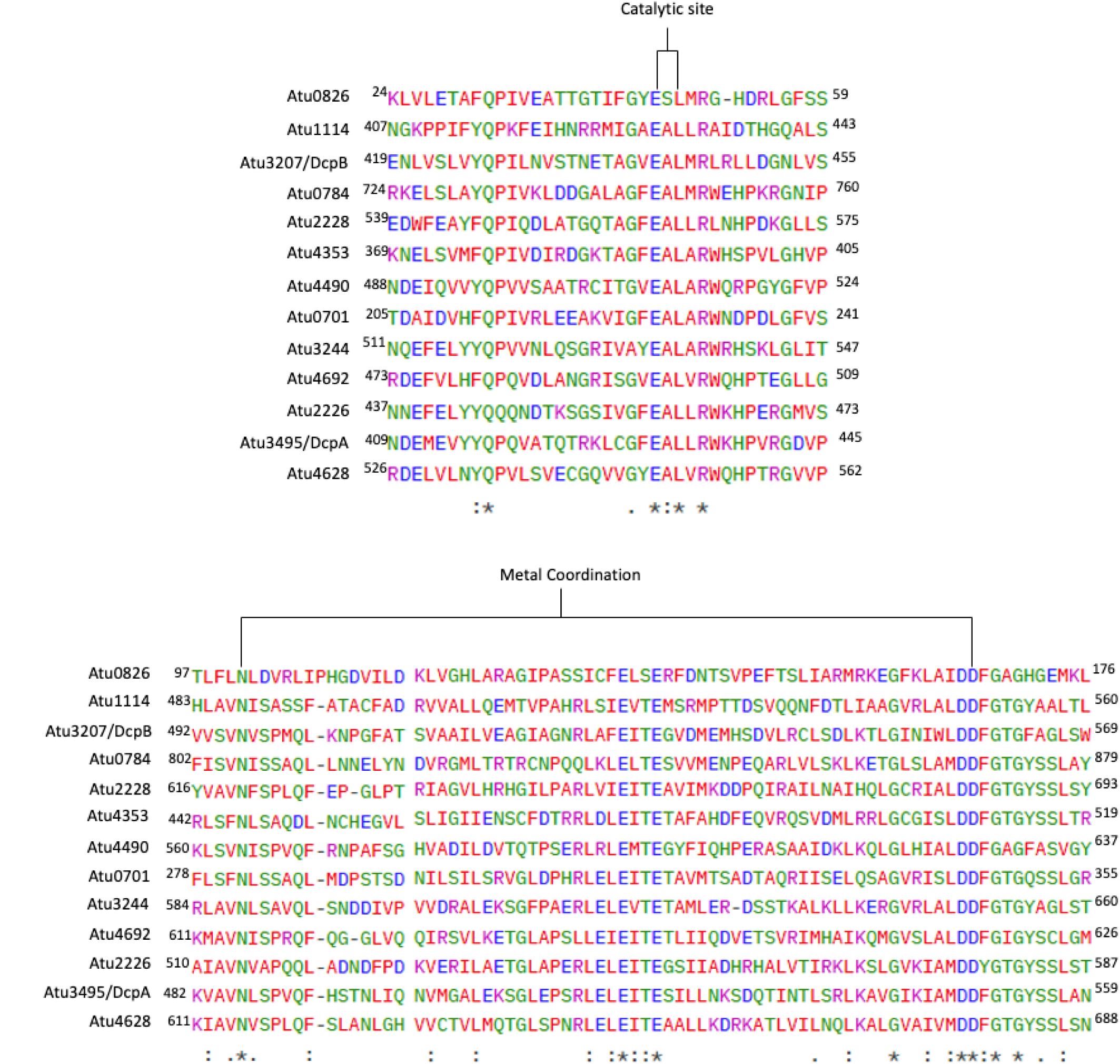
Dual-domain diguanylate cyclase/phosphodiesterase proteins from *A. tumefaciens* showed conserved catalytic site and metal coordination. Sequence alignment of the 13 predicted dual-domain proteins highlighting conserved elements of the phosphodiesterase domain. The EAL catalytic site and metal coordination are shown in the alignment. Atu0826 was used as the reference sequence. Protein sequences were aligned with Clustal Omega EMBL-EBI.

